# Commensal Microbiota Regulates Skin Barrier Function And Repair Via Signaling Through The Aryl Hydrocarbon Receptor

**DOI:** 10.1101/2020.12.05.413096

**Authors:** Aayushi Uberoi, Casey Bartow-McKenney, Qi Zheng, Laurice Flowers, Amy Campbell, Simon A.B. Knight, Neal Chan, Monica Wei, Victoria Lovins, Julia Bugayev, Joseph Horwinski, Charles Bradley, Jason Meyer, Debra Crumrine, Carrie Hayes Sutter, Peter Elias, Elizabeth Mauldin, Thomas R. Sutter, Elizabeth A. Grice

## Abstract

The epidermis forms a barrier that defends the body from desiccation and entry of harmful substances, while sensing and integrating environmental signals. The tightly orchestrated cellular changes required for the proper formation and maintenance of this epidermal barrier occur in the context of the skin microbiome. Using germ free mice, we demonstrate the microbiota is necessary for proper differentiation and repair of the epidermal barrier. These effects were mediated by the aryl hydrocarbon receptor (AHR) in keratinocytes, a xenobiotic receptor also implicated in epidermal differentiation. Murine skin lacking keratinocyte AHR was more susceptible to barrier damage and infection, during steady state and epicutaneous sensitization. Colonization with a defined consortium of human skin isolates restored barrier competence in an AHR-dependent manner. We reveal a fundamental mechanism whereby the microbiota regulates skin barrier formation and repair, with far-reaching implications for the numerous skin disorders characterized by epidermal barrier dysfunction.

## MAIN TEXT

## INTRODUCTION

The skin is the primary barrier between the human body and the environment and functions to prevent desiccation and entry of foreign and/or harmful substances. The barrier properties of the skin reside in the epidermis, a semi-permeable stratified epithelium that is formed as a result of keratinocyte terminal differentiation. Though continuously exposed to xenobiotic toxins, physical insults, and pathogenic microbes, the epidermis is also associated with diverse commensal microbial communities that are critical players in regulating skin physiology [1]. These microbial communities, collectively referred to as skin microbiota, are specialized to thrive in the unique nutrient and environmental conditions of this organ. The skin microbiome is topographically diverse, temporally complex, and distinct from other organs [2, 3]. How the commensal microbiota influences development of skin’s barrier function is undefined, as are the molecular mechanisms that mediate these interactions.

The barrier function of the skin may be conceptualized as four intertwined “levels” consisting of microbial, immune, chemical, and physical barriers [4]. The skin microbiome itself provides a barrier to pathogenic micro-organisms via a variety of different mechanisms e.g. production of proteases, antimicrobial peptides and antibiotics, and interference with quorum sensing [5]. This outermost microbial barrier also interacts with and mediates other functional levels of the cutaneous barrier. Skin microbiota play a fundamental role in the induction, training, and function of the skin immune barrier in part through the release of antimicrobial peptides, short-chain fatty acids, and polyamines [6]. Neonatal colonization by microbiota has long-lasting impacts on adult immune barrier as commensal skin microbes’ prime immune cells to differentiate between commensal versus pathogenic bacterium [7]. Bacterial lipases can hydrolyze lipids resulting in production of free fatty acids that impact the acidic surface pH of the skin, which dictates the chemical barrier of the skin [8, 9]. While studies in gnotobiotic mice suggest that epidermal differentiation and barrier genes are microbially regulated [10], mechanistic roles for the skin microbiota in development, regeneration, and function of the physical barrier are not well defined.

The epidermal permeability barrier (EPB) comprises of the stratum corneum and a complex system of tight junctions and adhesion complexes and their associated cytoskeletal networks that mediate cell-cell adhesion to create a mechanical barrier between the environment and underlying tissue [11]. Actively dividing keratinocytes in the stratum basale commit to terminal differentiation and move progressively into suprabasal layers, i.e. stratum spinosum, the stratum granulosum and eventually the stratum corneum [12]. In the stratum corneum, keratinocytes become flattened and denucleated (which are then called corneocytes), and plasma membranes are replaced with cornified envelopes. Lamellar bodies secrete their lipid-rich contents into the intercellular space between the corneocytes and are subsequently processed into barrier-providing lipid lamellae. Altogether, corneocytes, lipids and a complex network of trans-membrane proteins, provide a highly hydrophobic EPB against the environment. Microbial influences on this process of epidermal differentiation and EPB formation are not well understood, nor are the mechanisms whereby the EPB senses and responds to changes in the microbiota.

The sensing of xenobiotics, or compounds foreign to a living organism, is critical for barrier defense and homeostasis in the skin [13]. Keratinocytes function as sentinels that sense and respond to external stimuli [14]. Activation of xenobiotic receptors in keratinocytes induces expression of detoxification enzymes and membrane transporters that promote elimination of toxic compounds[15]. Accumulating evidence suggests that roles for xenobiotic receptors extend to cellular processes beyond xenobiotic metabolism that include cellular proliferation, tissue repair, and immune responses [13]. Microbes produce a plethora of small molecules and secondary metabolites, which are hypothesized to mediate their interactions with the host toward a mutualistic relationship [16]. The diversity of molecular signals produced by skin microbes, and how keratinocytes decipher and respond to them, remain largely unexplored.

The aryl hydrocarbon receptor (AHR) is a xenobiotic receptor that has emerged as a critical player in EPB development, function and integrity. Activation of the AHR, a ligand-activated transcription factor of the basic, helix-loop-helix motif-containing Per-ARNT-Sim family [17], induces a variety of epidermal differentiation and barrier genes, accelerates terminal differentiation, and increases stratum corneum thickness [18-22]. The AHR can be activated by halogenated and non-halogenated aromatic hydrocarbons, including dioxins such as 2,3,7,8-Tetrachlorodibenzo-p-dioxin (TCDD), polychlorinated biphenyls (PCBs) and polycyclic aromatic hydrocarbons (PAHs); clinically used drugs, food-derived molecules, endobiotics, and bacterial metabolites [23-26]. Microbial regulation of the AHR in the context of the skin barrier remains poorly understood, as well as the consequences of perturbing the commensal microbiota with respect to EPB function and defense.

In addition to congenital barrier deficiencies, epidermal barrier dysfunction is a hallmark of inflammatory skin diseases, including atopic dermatitis and psoriasis, and predisposes skin to infections [27-29]. Additionally, epicutaneous sensitization, as a result of epidermal barrier dysfunction, may lead to atopic and allergic disease [30].Thus, there is an urgent scientific and clinical need to define the mechanistic basis by which the commensal microbiota regulate homeostatic barrier function, as such mechanisms provide new targets for prevention and/or intervention in skin barrier deficiencies.

Here, we investigated the role of commensal microbiota in regulation of permeability barrier homeostasis of skin. We found that commensal microbes are necessary for normal epidermal differentiation, EPB function, and repair. These effects were mediated by microbial signaling through the keratinocyte AHR. Murine skin lacking keratinocyte AHR signaling displayed increased barrier permeability, enhanced susceptibility to infection by *S. aureus*, and increased pathology in a model of atopic dermatitis. We show that topical colonization with a defined consortium of human skin commensals improves EPB function in murine germ-free skin and models of barrier dysfunction. Our findings reveal a fundamental role for the commensal skin microbiota in regulating the physical integrity and repair of the skin barrier, provides mechanistic insights into microbial-skin crosstalk, and uncovers therapeutic targets for improving skin barrier function.

## RESULTS

### Epithelial development and differentiation programs are impaired in germ free skin

To characterize microbially-mediated regulation of homeostatic epithelial gene expression programs, we performed RNA-seq on epidermal sheets isolated from dorsal skin of *C57BL/6* mice of 3 different colonization states (n=8 mice each. **Figure 1A**): Specific pathogen free (SPF) mice that were conventionally raised in presence of microbiota, germ free (GF) mice born and raised in sterile gnotobiotic isolators, and a third group of mice that were born GF and then colonized (COL) with SPF microbiota for 2 weeks. Within 2 weeks, COL mice were colonized with microbiota from SPF mice as validated by 16S ribosomal RNA (rRNA) gene sequencing (**Figure 1B**). We identified differentially expressed genes (DEGs) between colonization states by training negative-binomial linear models using DESeq2 R package and 3-way comparisons. This analysis revealed 6396, 427, and 3232 DEGs for SPF vs. GF, COL vs. GF and SPF vs. COL comparisons, respectively (**Figure 1C, Table S1**). We focused on the 396 shared DEGs of SPF and COL epidermis when compared to GF epidermis (**Figure 1D**). We reasoned that this subset of DEGs meet the criteria of being induced and sustained by microbial colonization, suggesting homeostatic control. The 396 DEGs were significantly enriched for biological functions such as skin development, keratinocyte differentiation, and epidermis development (**Figure 1E**). This result suggests the microbiota plays an important role in epithelial barrier formation.

**Figure 1.**
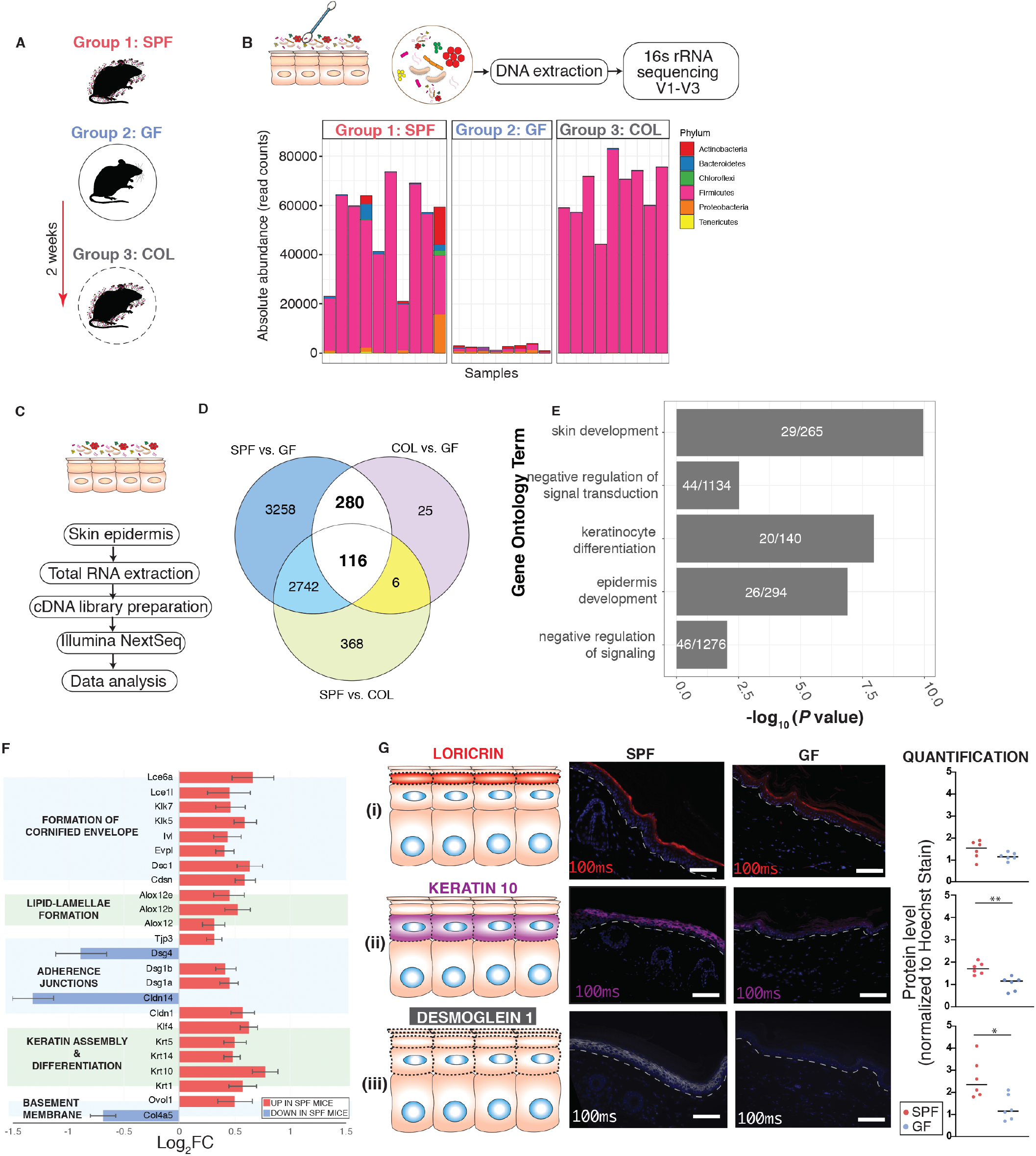
Commensal microbiota regulates epithelial barrier genes. **(A)** Three groups of mice were employed, specific pathogen free (SPF), germ free (GF), and germ-free mice colonized (COL) with SPF microbiota for 2 weeks. **(B)** Skin microbiota composition determined by 16S rRNA gene sequencing. Y-axis indicates absolute read counts of most abundant phylum (by relative abundance in the dataset) for each mouse (x-axis). **(C)** RNA-seq workflow. **(D)** Overlap of differentially expressed genes when comparing groups of gnotobiotic mice. **(E)** Shown in white are the number of genes that were further analyzed for uniquely enriched gene ontology biological process terms for aforementioned DEGs. Shown on the y-axis are the uniquely enriched terms, with *p*-values indicated on the x-axis. *P*-values are based on Fisher’s exact test and FDR-adjusted under dependency using the “BY” method. **(F)** To examine DEGs involved in epithelial barrier function, we manually curated genes involved in different facets of epithelial barrier: keratinization, cornified envelope formation, adherence and gap junction, basement membrane function, barrier development, differentiation and intercellular lipid-lamellae processing. Shown here is a snapshot of key genes that were differentially expressed in the SPF vs GF subset (*p<0*.*001*). Horizontal bars represent the Log2 fold-change comparison (genes upregulated in SPF: log2FC> 0, downregulated in SPF: log2FC < 0). Error bars represent standard error estimate for the log2 fold-change computed using the DESeq2 package. **(G)** Tail-skin from SPF and GF mice. Immunofluorescence-based detection of differentiation markers (i) loricrin (red) and (ii) keratin-10 (purple) and adhesion marker (iii) desmoglein-1(grey). Nuclei are counter-stained with Hoechst stain (blue). Images were taken at constant light exposure (100ms for proteins of interest and 10ms for DAPI channel) and then overlaid for representation. White dashed-line indicates boundary separating epithelial-stromal compartments. Scale bar (10µm) is indicated in white (bottom-right). For quantification 10-12 random images were taken in a blinded fashion for each mouse tissue (n=6) at constant light exposure and processed through ImageJ. For each image integrated density of signal was normalized to Hoechst stain signal from the same area. Each dot corresponds to average normalized signal across 10-12 images for each mouse. Asterisk indicates statistical significance (p<0.05, T test, two-sided). See also Table S1, S2, S3, S4 and Figure S1.

To further examine DEGs involved in epithelial barrier function, we manually curated genes involved in different facets of epithelial barrier: keratinization, cornified envelope formation, adherence and gap junction, basement membrane function, barrier development, differentiation and intercellular lipid-lamellae processing (**Table S2)**. Focusing on the SPF vs. GF subset of DEGs, multiple genes across each of these categories were expressed at lower levels in GF mice (**Figure 1F, Tables S3, S4**). In particular, genes critical for cornified envelope formation [e.g. *involucrin (Ivl), envoplakin (Evpl)*] and its desquamation, [e.g. *Kallikrien-related peptidases 5, 7(Klk5, 7)*] were downregulated in GF skin. We hypothesized that such significant and widespread differences in gene expression would result in structural differences between GF and SPF skin. However, consistent with prior reports [10, 31] we did not notice any overt differences between the epithelial organization of GF vs SPF mice by traditional histopathological examination (**Figure S1**). Analysis of skin ultrastructure by electron microscopy showed that overall, SPF mice had a greater number of individual layers within the stratum corneum than GF mice (**Figure S1**). Immunofluorescence-based analysis of molecular biomarkers showed decreased expression of loricrin and cytokeratin-10 in skin of GF mice (**Figure 1G**) that are implicated in barrier integrity [32-34]. Among genes that were downregulated in GF skin were tight and adherens junctions family members (**Figure 1F**) such as *tight junction protein 3 (Tjp3), desmogleins 1a-b (Dsg1)*, and *claudin-1 (Cldn1)* that are critical players involved in skin barrier formation (reviewed in [35]). Remarkably, tight junction integrity appeared compromised in the suprabasal epithelium as evident by downregulated and diffused expression of *Dsg1* in GF mouse epidermis (**Figure 1G)**. Disruption of *Dsg1* is associated with improper formation of desmosomes in suprabasal epithelia and has been associated with skin barrier impairment [36-38]. Together, these findings suggest the hypothesis that skin barrier formation requires the commensal microbiota.

### Commensal microbiota promotes skin barrier function and repair

A fully functioning stratum corneum closely controls the water concentration gradient in the skin such that passive diffusion of water occurs from inner layers towards the outside. Barrier disruption compromises the ability of the stratum corneum to maintain this water concentration gradient and results in increased transepidermal water loss (TEWL), measured using a sensor for water vapor flow density [39, 40]. Low TEWL values are indicative of intact skin and increased TEWL is associated with a disrupted barrier (**Figure 2A, Panel i**). Under basal conditions, GF mice had slightly increased TEWL compared to SPF mice corroborating the findings from ultrastructure analysis (**Figure S2**). Skin barrier of GF mice was perturbed more readily with tape stripping than SPF skin (**Figure S2**), consistent with fewer layers of stratum corneum in GF mice. After comparable insults following tape-strip injury (TEWL 15-20 g/m^2^/h), SPF mice more rapidly repaired their barrier compared to GF mice when measured over a period of 24 hours (**Figure 2B**). We observed similar delays in barrier recovery in GF *Rag1*^*−/−*^ mice (that lack mature T and B-cells) compared to age-matched SPF *Rag1*^*−/−*^ mice (**Figure 2C, Figure S2**) suggesting that microbially-regulated adaptive immune responses are not responsible for the delayed barrier repair phenotype. Skin from GF *Rag1*^*−/−*^ mice also showed decreased expression of genes involved in terminal differentiation and formation of transmembrane junctions compared to SPF *Rag1*^*−/−*^ mice (**Figure S2)** as seen in wild-type mice.

**Figure 2.**
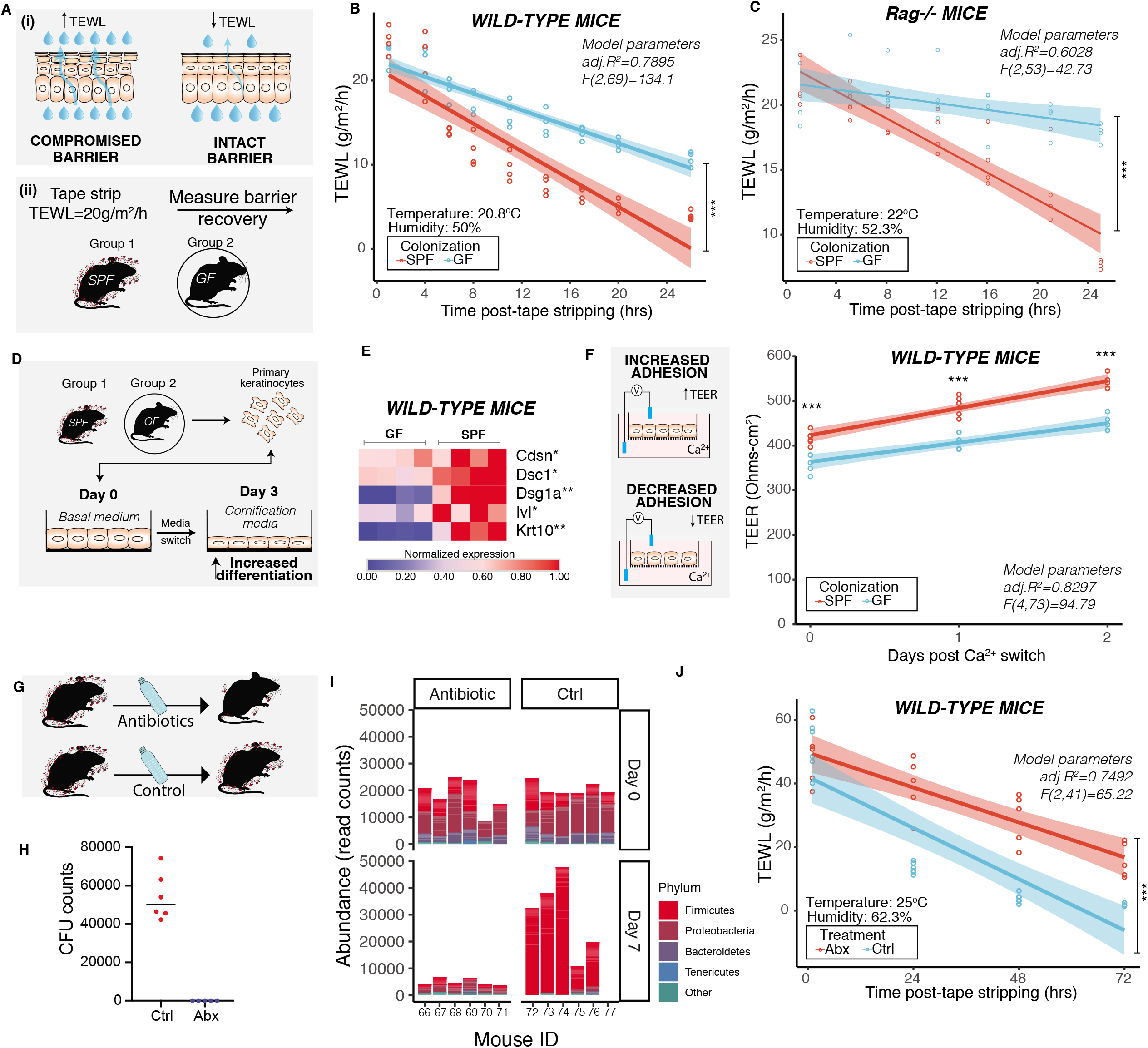
Commensal microbiota promotes skin barrier repair function. **(A)** Schematic depicts (i) principle of measuring transepidermal water loss (TEWL) to assess barrier repair function in adult mice (6-8 weeks old). (ii) Experimental design for assessing barrier recovery. Dorsal skin of mice was tape-stripped to achieve comparable insults and TEWL was measured up to 24 hours post-tape stripping. Effect of colonization of microbes was assessed by comparing age-matched germ-free (GF) and specific pathogen-free (SPF) mice (n=4 mice per group) in **(B)** wild-type C57/BL6 mice [ANCOVA, *F (1,69) =50*.*649, ***P<0*.*001)*] and **(C)** *Rag1*^*−/−*^ mice [ANCOVA, *F (1,53) =188*.*1, ***P<0*.*001)*]. **(D)** Primary mouse keratinocytes were derived wild-type GF (n=4) and SPF (n=4) C57/BL6 mice and grown in 5% FBS and 1.6mM Ca^2+^ for three days to induce terminal differentiation. **(E)** Expression of genes involved in differentiation [lnvolucrin *(Ivl)*, cytokeratin-10 *(Krt10)*] and adherence [Corneodesmosin *(Cdsn)*, Desmocollin-1 *(Dsc1)*, Desmoglein-1a *(Dsg1a)*] was assessed by qRT-PCR. Cycle thresholds were normalized to housekeeping genes (*Rplp2, Sptbn1* and *18S rRNA*) and normalized relative to Cq values of SPF condition. Each square represents average readings from keratinocytes (n= 4 technical replicates) derived from an individual mouse (n=4 mice per group). \**P<0*.*01*, ** *P<0*.*001* by T-test adjusted by Bonferroni correction. **(F)** Primary keratinocytes were grown on transwells in 5% FBS and 1.6mM Ca^2+^. Epithelial adhesion was assessed by measuring transepithelial electrical resistance (TEER) at indicated time points. Data from one experiment is represented for visualization (See Figure S2). One dot represents average TEER readings from technical replicates (n=3) derived from one individual mouse (n=4 mice per group). \*\*\**P<*0.001 by two-way ANOVA adjusted for multiple experiments. **(G)** To decrease skin microbial burden, wild-type SPF mice were treated with antibiotic cocktail (n=5) or vehicle (n=6) for two weeks. **(H)** To determine microbial burden, mice were swabbed 14 days after treatment and colony forming units (CFU) were determined. **(I)** Genomic DNA was extracted from swabs collected at baseline (Day-0) and after one week of treatment (Day 7). V1-V3 region was amplified and analyzed by 16S rRNA gene sequencing and overall abundance i.e. total operational taxonomic unit (OTUs) counts belonging to different phyla in each sample are depicted. Phyla with total read count <1000 are grouped into ‘Other’. (See Figure S3) **(J)** At the end of two weeks the two groups of mice were tape stripped to achieve comparable insults and TEWL was measured and plotted against time [ANCOVA, *F (1,41) =26*.*315, ***P<0*.*001)*]. TEWL/TEER vs time readings were fitted by linear modeling (in **B, C, F** and **J**) and significance was assessed by ANCOVA analysis. Modeling parameters (adjusted *R*^*2*^ and *F*-statistics) are indicated on top-right for each plot. Span indicated by shaded area represents 95% CI. Temperature and humidity conditions during TEWL measurement are indicated for each experiment. Also see Figures S2 and S3.

Basal keratinocytes undergo a spatiotemporal and highly controlled differentiation program dependent on intracellular calcium flux to establish and maintain barrier [41]. In presence of high calcium, primary epidermal keratinocytes can differentiate *in vitro* to express genes involved in formation of cornified envelope [42]. We derived murine primary epidermal keratinocytes from GF and SPF skin, respectively, and exposed them to high calcium containing medium (**Figure 2D**). Expression of genes involved in terminal differentiation [*cytokeratin-10 (Krt10), involucrin (Ivl)*] and formation of transmembrane junctions [*Corneodesmosin (Cdsn), Desmocollin-1(Dsc1), Desmoglein-1a (Dsg1a)*] were reduced in GF keratinocytes compared to SPF keratinocytes (**Figure 2E, Figure S2)**. Additionally, GF keratinocytes had decreased transepithelial electrical resistance (TEER) which is indicative of decreased transmembrane junction strength (**Figure 2F, Figure S2**).

To further examine the implications of perturbing the microbiota on skin barrier function we used an antibiotic depletion model (**Figure 2G**). Prior studies had shown that antibiotics traditionally used to disrupt gut microbiota in mice were not sufficient to disrupt skin microbiota in mice [43]. We developed a new regimen consisting of antibiotics (Metronidazole, Sulfamethoxazole, Trimethoprim, Cephalexin and Enrofloxacin) that are administered orally in hospitals and veterinary clinics to target skin bacteria [44-47] and were able to inhibit prominent murine skin commensal *Staphylococcus xylosus* [48, 49]. Oral administration of antibiotics for two weeks diminished microbial burden on skin as observed by both quantitative cultures and 16S rRNA gene sequencing (**Figure 2H, I**) but did not significantly affect microbial burden in the gut (**Figure S3)**. Antibiotic-treated mice were delayed in barrier repair compared to control mice that were treated with vehicle (**Figure 2J**). Together, these data confirm a role for commensal microbes in promoting skin barrier function and repair.

### Aryl hydrocarbon receptor pathway is attenuated in germ free skin

The sensing of external physiological and chemical signals is critical for barrier defense and homeostasis in the skin [13]. Therefore, we hypothesized that xenobiotic receptors that act as epithelial sensors and relay microbial signals would be among those genetic pathways dysregulated in GF epidermis. Previous studies have identified at least 304 xenobiotic processing genes (XPGs) in mice, which encode the enzymes, transporters, and transcription factors required to metabolize xenobiotics [50]. We found that 52/304 XPGs were differentially expressed (*P<0*.05) in skin of SPF and GF mice, and the majority were upregulated in SPF mice (n=43/52; **Table S5**). The pregnane X receptor (PXR, *NR1I2*), constitutive androstane receptor (CAR, *NR1I3*), peroxisome proliferator-activated receptor-alpha (PPARa) and aryl hydrocarbon receptor (AHR) are key transcription factors that regulate xenobiotic processing in skin [51]. Of these, only the AHR gene was differentially expressed and was upregulated in SPF skin compared to GF skin. Canonically, after ligand binding, the AHR translocates to the cell nucleus and binds DNA at xenobiotic responsive elements (XRE), to regulate transcription of target genes [52]. Expression of key downstream target genes, i.e. cytochrome-p450 *Cyp1a1* and molecular chaperones *Hsp90aa1* and *Hsp90ab1* that respond to AHR activation, were also downregulated in GF murine epidermis (**Figure 3A**). Primary keratinocytes derived from GF skin were also impaired in expression of these genes (**Figure S2**). Overall, microbiota-mediated upregulation of *AHR* was consistent with changes in its canonical pathway, suggesting that regulation of xenobiotic processing genes in the skin may be mediated through the *AHR*.

**Figure 3.**
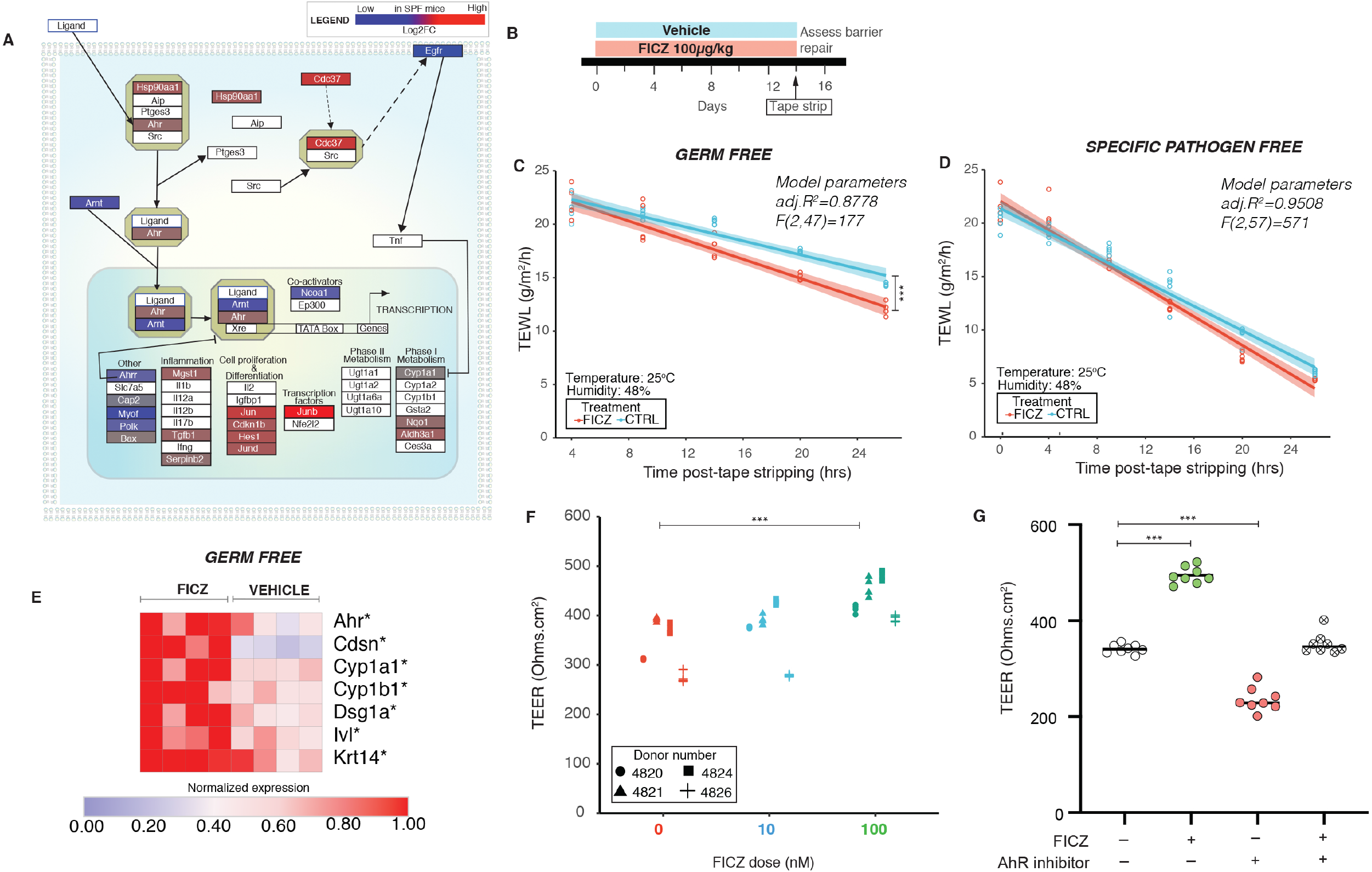
Activation of aryl hydrocarbon receptor (AHR) signaling in skin rescues barrier dysfunction in germ free mice. **(A)** Differentially expressed genes (DEGs) in SPF vs GF mice, as identified by RNAseq, were mapped onto the AHR pathway. Log2FC of significant DEGs (*P<0*.*01)* in SPF mice are represented in colored boxes according to the legend. **(B)** Schematic illustrating experimental design. Age matched 6-week-old, GF and SPF (n=5 mice/group) C57/BL6 mice, respectively, were treated daily with 6-formylindolo[3,2-b] carbazole (FICZ) at 100µg/kg or vehicle for 2 weeks. At end of treatment dorsal skin of mice was tape-stripped to achieve comparable insults (∼20g/m^2^/h) and TEWL was measured up to 24 hours post-tape stripping to assess barrier recovery. TEWL vs time readings were fitted by linear modeling and covariance was assessed by ANCOVA. Barrier recovery was compared in **(C)** GF mice [*F (1,47) =21*.*9, ***P<0*.*001)*] and **(D)** SPF mice *[F(1,57)=2*.*98, *P=0*.*0492]* that were either treated with FICZ or vehicle. **(E)** Expression of genes [*Ahr, Cdsn, Cyp1a1, Cyp1b1, Dsg1a, Ivl* and Krt14] was assessed by qRT-PCR in GF mouse skin treated with FICZ or vehicle (4 mice per group). Cycle thresholds were normalized to housekeeping genes (*Rplp2, Sptbn1* and *18s rRNA*) and normalized relative to Cq values of FICZ treatment. \**P<0*.*05*, by T-test adjusted by Bonferroni correction. **(F)** Primary human keratinocytes grown on transwells (in 5% FBS and 1.6mM Ca^2+^) in presence of FICZ (0nM, 10nM and 100nM) for three days and transepithelial electrical resistance (TEER) was measured. Cells from different donors are represented by different symbol. See Figure S4. **(G)** Primary human keratinocytes (in 5% FBS and 1.6mM Ca^2+^) were treated as indicated with FICZ and/or AHR inhibitor at 100nM doses. TEER values at the end of three days of treatment are reported. \*\*\**P<0*.*001* by T-test for panels F and G.

Since the AHR can activate multiple signaling pathways [53], we explored the impact of microbial colonization on expression of genes reported to be a part of canonical and non-canonical AHR signaling networks (**Figure 3A**). In this context, we saw upregulation of several genes in SPF mice, including those involved in cell proliferation, differentiation, and inflammation (e.g. *Hes1, Jun* and *Tgfb1)*. Together these findings suggest the AHR as a potential mechanism by which skin microbes modulate epithelial barrier integrity.

### Treatment with an AHR agonist improves barrier function and recovery in germ free mice

We hypothesized that if the GF skin barrier phenotype we observed was due to attenuated AHR signaling, then activation via AHR ligand would improve barrier recovery. We treated adult GF mice topically with AHR ligand 6-Formylindolo[3,2-b] carbazole (FICZ), daily for 2 weeks at a low dose (100µg/kg) (**Figure 3B**), a regimen shown to induce expression of *Cyp1a1* [54]. FICZ is a tryptophan photoproduct and is a well-characterized AHR ligand in skin [55, 56]. At the end of treatment, we compared the rate of barrier recovery in tape-stripped GF mice that were either treated with FICZ or vehicle (**Figure 3C**). We observed that FICZ significantly accelerated barrier recovery in GF mice. In parallel, we also treated SPF mice with FICZ at the same dose and observed that FICZ accelerated early stage barrier recovery (**Figure3D**). FICZ treatment activated AHR signaling in the treated group as *CYP1A1* expression was induced in all FICZ treated mice in comparison to the untreated group (**Figure 3E**). Additionally, FICZ treatment increased expression of genes implicated in barrier repair (**Figure 3E**) in GF mice. AHR upregulation increases expression of genes involved in epidermal differentiation and formation of gap junctions [57]. In primary human keratinocytes, FICZ increased epithelial resistance as measured by TEER (**Figure 3F, Figure S4**), which was diminished by co-treatment with an AHR inhibitor (**Figure 3G**). Overall, activation of the AHR in GF skin improves epidermal barrier recovery, supporting a mechanistic role for the commensal microbiota in homeostatic regulation of the AHR.

### Mice deficient in epithelial AHR have a defective skin barrier and are more susceptible to infection

AHR is expressed in a variety of cell types in the skin, but abrogating AHR function in keratinocytes has been suggested to impact skin barrier [58]. To confirm the role of keratinocyte AHR in skin barrier function, we generated mice where the *floxed Ahr* allele (*Ahr*^*f/f*^) was conditionally knocked out in the skin epithelia using a *Cre* driven by the keratin-14 promoter (*K14*^*Cre/+*^*Ahr*^*f/f*^). While littermates that retained AHR function (*Ahr*^*f/f*^) repaired their barrier within 24 hours following tape-strip disruption, barrier repair was significantly diminished in *K14*^*Cre/+*^*Ahr*^*f/f*^ skin (**Figure 4A**). Additionally, keratinocytes derived from *K14*^*Cre/+*^*Ahr*^*f/f*^ mice showed increased TEER compared to *Ahr*^*f/f*^ keratinocytes (**Figure 4B**).

**Figure 4.**
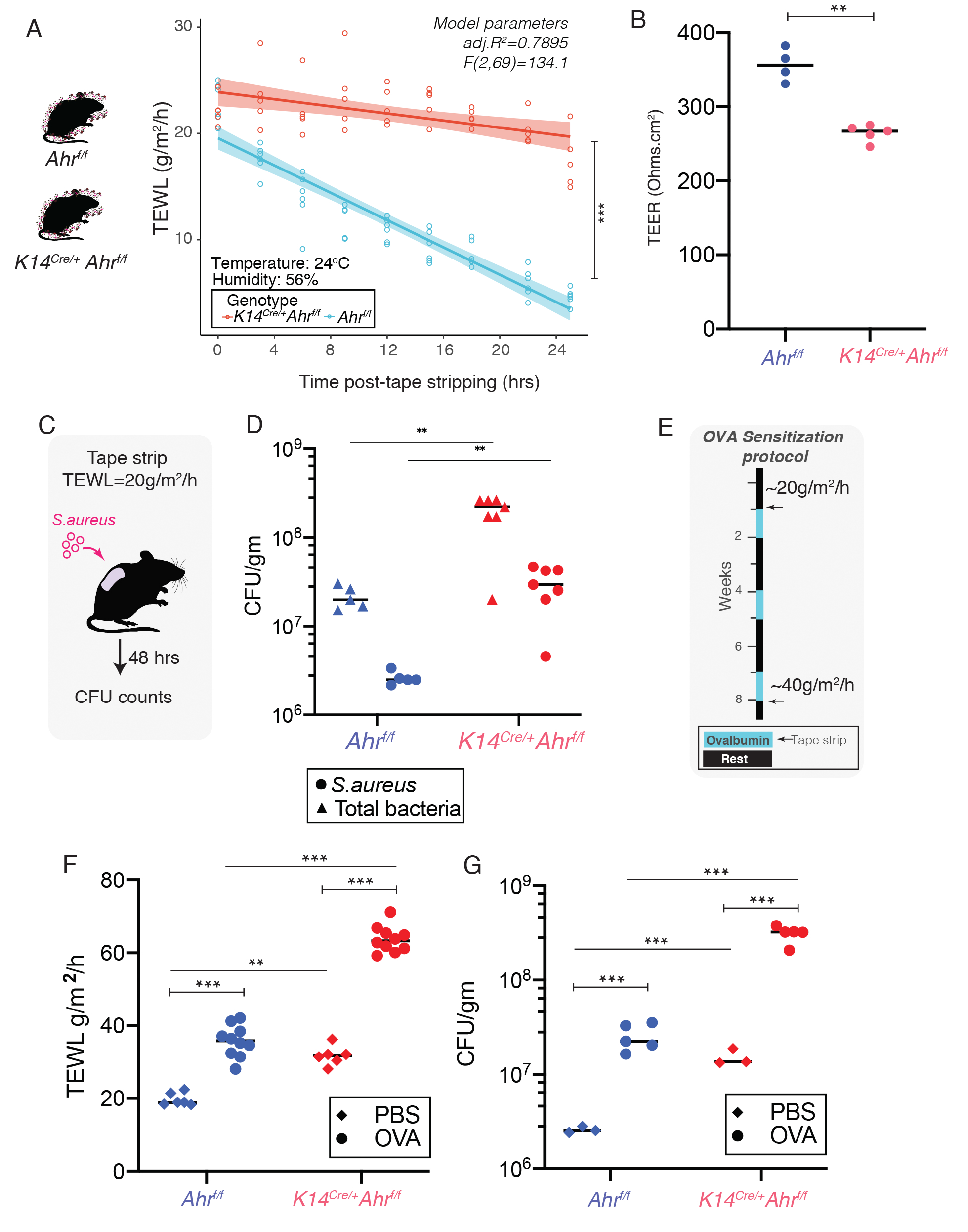
Loss of aryl hydrocarbon receptor (AHR) in keratinocytes impairs skin barrier in mice. **(A)** *Ahr* floxed *(Ahr*^*f/f*^) allele was knocked out in mice using Cre driven by a keratin-14 promoter *(K14*^*Cre/+*^*Ahr*^*f/f*^). Dorsal skin was tape-stripped to achieve comparable insults (20-25 g/m^2^/h) and TEWL recovery curves were compared between *Ahr*^*f/f*^ (n=5) and *K14*^*Cre/+*^ *Ahr*^*f/f*^ (n=6) mice [ANCOVA, *F (1,96) =131*.*34, ***P<0*.*001)*]. **(B)** Primary mouse keratinocytes were derived from mice and polarized in 5% FBS and 1.6mM Ca^2+^ for three days and transepithelial electrical resistance (TEER) was measured (\*\**P<0*.*005*, T-test). **(C)** *Ahr*^*f/f*^ (n=5) and *K14*^*Cre/+*^ *Ahr*^*f/f*^ (n=7) mice were tape-stripped (TEWL=20 g/m^2^/h) and 10^7^ CFU *S. aureus* containing tdTomato was applied to back skin. **(D)** 48 hours post-infection tissue was collected, weighed, homogenized and plated. *S. aureus* (visible as red colored colonies) and total bacterial colonies were counted (\*\**P<0*.*005*, T-test). **(E)** Model for atopic dermatitis induced by repeated epicutaneous sensitization of tape-stripped skin with ovalbumin (OVA) or vehicle (PBS) [30] was implemented. Mice were tape stripped at the beginning of the experiment (TEWL∼20 g/m^2^/h) and OVA was applied daily for 7 days, 3 times with rest for 2 weeks between each treatment. At the end of final treatment, mice were tape-stripped (TEWL∼40 g/m^2^/h) and 24 hours later **(F)** TEWL levels were assessed and **(G)** *S. aureus* was applied to back skin, *S. aureus* CFUs were determined (as described in E). Statistical significance in panels G and H were assessed using a 2-way ANOVA (*\*\*P<0*.*005*, \*\*\**P<0*.*0005*). (See Figure S4)

Patients with diseases of skin barrier impairment such as atopic dermatitis (AD) are highly susceptible to colonization and infection by pathogens including *S. aureus* [59]. To test if AHR-dependent barrier impairment leads to increased infection, we topically applied *S. aureus* to tape-stripped skin [60] and quantified bacteria following 48 hours. We observed enhanced infection of *S. aureus* and increased overall bacterial burden on tape stripped skin (**Figure 4C**) of *K14*^*Cre/+*^*Ahr*^*f/f*^ mice compared to AHR sufficient controls (**Figure 4D**). Since barrier dysfunction is a hallmark in the development of AD [61], and mice lacking AHR are impaired in barrier repair, we hypothesized that these mice will be more prone to barrier damage, infection, and atopic disease. We adapted a mouse model of AD (**Figure 4E**) induced by repeated epicutaneous sensitization of tape-stripped skin with ovalbumin (OVA) [30]. In this model, *K14*^*Cre/+*^*Ahr*^*f/f*^ skin was exacerbated in disease pathology, with enhanced TEWL (**Figure 4F**) and increased susceptibility to *S. aureus* infection compared to *Ahr*^*f/f*^ mice (**Figure 4G**). Together, these data demonstrate that AHR function in keratinocytes is essential for barrier function, and loss of AHR can lead to enhanced barrier damage in the setting of inflammatory skin disease and facilitate bacterial entry.

### Human skin microbial consortium restores barrier repair and function via the AHR

In the gastrointestinal tract, the AHR is activated by the gut microbiota [62] to enhance intestinal barrier integrity by inducing tight junction proteins in intestinal epithelial cells [63]. However, the relation between skin commensals and cutaneous AHR-dependent barrier regulation is unexplored. Given our observations that the commensal microbiota directly impacts barrier repair (**Figure 1, 2**) and regulates the AHR genetic pathway (**Figure 3**) in murine skin, we hypothesized that topical association of GF skin with human skin commensals would activate the AHR and restore skin barrier repair. To test this hypothesis, we first curated a collection of cultured skin microbes that were abundant and prevalent on healthy human skin. Referred to as Flowers’ Flora, the collection consists of members of Firmicutes phylum i.e. *Staphylococcus epidermidis, S. warneri, S. hemolyticus* and members of Actinobacteria phylum i.e. *Micrococcus luteus, Corynebacterium aurimucosum* (**Figure 5A**). These skin microbes activated AHR in keratinocytes as determined by way of a reporter assay consisting of the AHR reporter element conjugated with *Cyp1a1* (**Figure 5B**). Flowers’ Flora colonized murine GF skin as determined by bacterial culture swabs and species-specific qPCR analysis (**Figure 5C, D**). After two weeks of colonizing with this defined consortium of human skin commensals (**Figure 5E**) barrier recovery function in GF skin was restored (**Figure 5F**). Skin of colonized mice showed elevated expression of differentiation genes as well as *Cyp1a1* (**Figure 5G**) compared to GF skin. Upon terminal differentiation, keratinocytes derived from colonized mice were enhanced in TEER compared to GF controls (**Figure 5H**). Colonization of *K14*^*Cre/+*^*Ahr*^*f/f*^ mice with Flowers’ Flora did not improve barrier repair (**Figure 5I**), further supporting an AHR-dependent mechanism. Finally, to determine if Flowers’ Flora could prevent barrier damage in AD-like disease, we pre-colonized wild-type SPF murine skin prior to short-term epicutaneous sensitization with OVA (**Figure 5J**). Pre-colonization with Flowers’ Flora significantly improved barrier recovery compared to skin that was not pre-colonized (**Figure 5K**). Together, these findings indicate that skin commensal microbes signal through the AHR to maintain homeostatic control of epidermal barrier integrity, and suggest new targets for preventing and/or treating epidermal barrier dysfunction.

**Figure 5.**
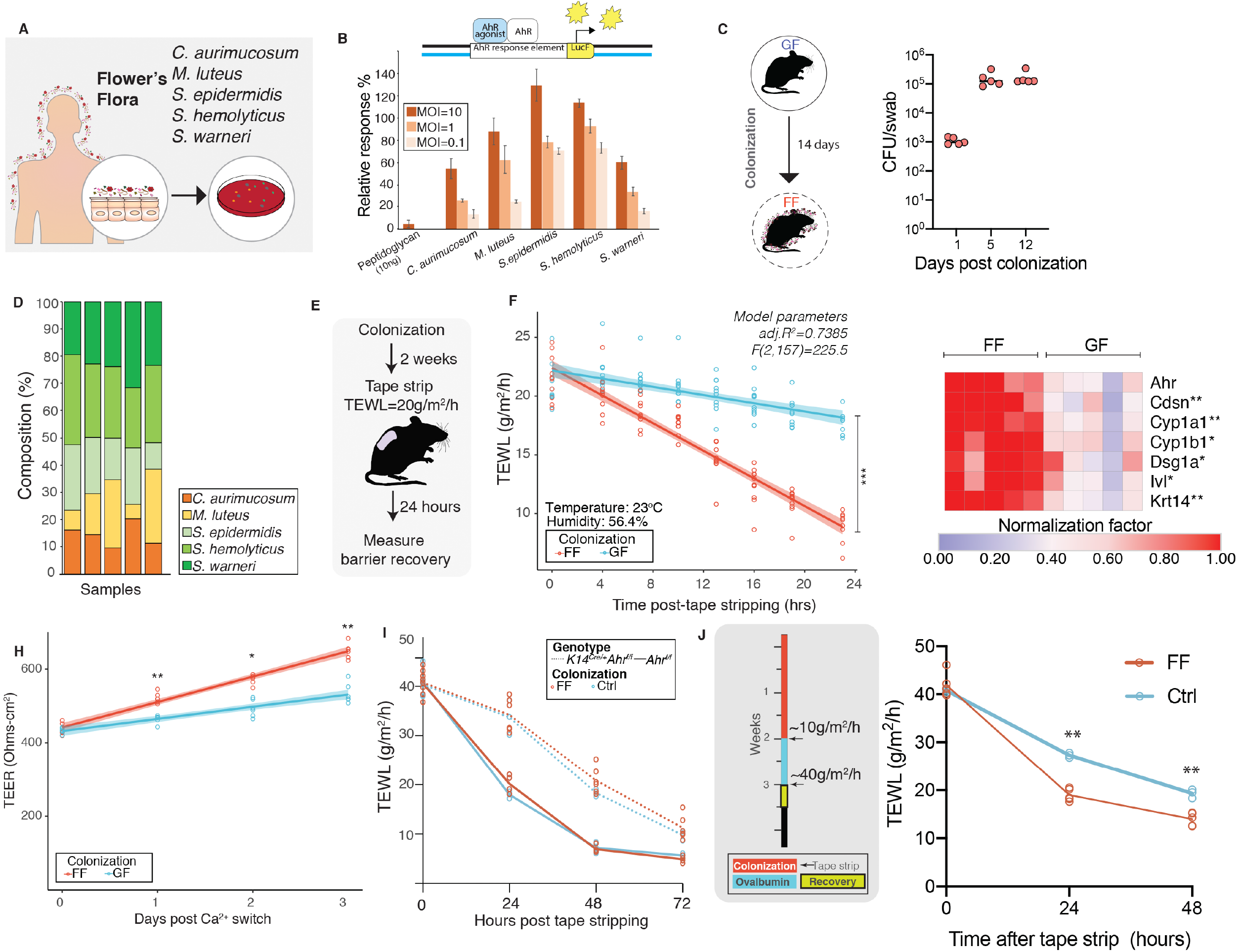
Commensal microbes curated from human skin restores skin barrier function in germ-free mice via AHR activation. **(A)** Curation of bacteria for Flowers’ Flora (FF) consortium. **(B)** HaCaT cells (10^6^ cells/well) were co-transfected with plasmids containing a *Firefly* luciferase reporter conjugated to *Cyp1a*-AHR response element and a *Renilla* luciferase as transfection control. Transfected cells were treated with indicated bacteria at indicated multiplicity of infection (MOI) and luminescence was measured. *Firefly* luciferase activities were normalized to *Renilla* luciferase levels and relative response compared to 10nM FICZ treatment (positive control) was computed. **(C)** Germ-free mice were colonized with FF daily for two weeks. Mice were swabbed at indicated days and CFUs were enumerated by plating on blood agar plates. **(D)** To determine whether individual bacteria of FF colonized skin, genomic DNA was extracted from skin swabs collected from mice at day 14 and qPCR analysis was conducted using species-specific primers for each bacterium and percentage composition relative to total 16S rRNA was determined. **(E)** Two-weeks post colonization mice (n=5 mice/group) that were either germ-free (GF) or colonized with FF were tape stripped (TEWL ∼20-30 g/m^2^/h) and **(F)** barrier recovery was assessed by TEWL (ANCOVA, *F(1,157)=181*.*25, P<0*.*0001)*. **(G)** Expression of genes [*Ahr, Cdsn, Cyp1a1, Cyp1b1, Dsg1a, Ivl* and Krt14] was assessed by qRT-PCR in mouse skin treated with FF or vehicle (GF). Cycle thresholds were normalized to housekeeping genes (*Rplp2, Sptbn1* and *18s rRNA*) and normalized relative to Cq values of FF treatment. \**P<0*.*05* and \*\**P<0*.*005* by T-test adjusted with Bonferroni correction. **(H)** Primary mouse keratinocytes were derived and polarized in 5% FBS and 1.6mM Ca^2+^ for three days and transepithelial electrical resistance (TEER) was measured (\*\**P<0*.*005*, T-test). **(I)** To test if improved barrier recovery via FF is through AHR, *K14*^*Cre/+*^*Ahr*^*f/f*^ (n=6) were pre-colonized as shown in Fig. 4F and compared to *K14*^*Cre/+*^*Ahr*^*f/f*^ (n=3) that were treated with Control (Ctrl) [ANCOVA, *F(1,61)=0*.*1191, P=0*.*73115*]. Additionally, *Ahr*^*f/f*^ mice that were colonized (n=4) and untreated (n=3) were included in comparisons. **(J)** To test if pre-colonization with FF could improve barrier recovery in an epicutaneous sensitization model, we pre-colonized C57BL6/J mice (n=5) with FF for 2-weeks. At the end of colonization, mice were tape-stripped and treated with Ovalbumin for 1-week. Mice were subjected to comparable insults (TEWL∼40g/m^2^/h) and barrier recovery kinetics were compared in FF colonized versus control (non-colonized) mice. FF colonized mice showed improved barrier recovery compared to control mice [ANOVA, *\*\*P<0*.*01*]. (See Figures S5).

## DISCUSSION

The skin microbiome provides the first level of barrier defense to the human body. While commensal skin microbes have demonstrated effects on immune and chemical barriers of the skin, their regulation of the physical barrier is not well-defined. Proper epithelial differentiation and cornification is essential for formation of the epithelial barrier [11, 64, 65]. By using comparative transcriptomics of gnotobiotic mice, we identified genes involved in skin development, differentiation and barrier function as top candidates for microbial regulation. Using multiple models of microbial depletion, loss of commensal microbiota impaired the barrier repair function of the skin. We mechanistically linked the xenobiotic receptor AHR in mediating microbial signals to the keratinocyte to boost epithelial differentiation and adhesion, and thus integrity of the EPB. The absence of such signaling resulted in exacerbated pathology and increased bacterial infection in models of barrier disruption and epicutaneous sensitization. Finally, we showed that topical association with a consortium of human skin commensals restores skin barrier function and integrity, an effect that was dependent on the keratinocyte AHR.

Our findings parallel those in the gastrointestinal tract, where gut commensals have been demonstrated to regulate intestinal barrier formation by modulating epithelial turnover [66] and controlling mucus production [67]. However, unlike the simple mucosal epithelium that provides the intestinal barrier, the skin is composed of a multi-layered stratified squamous epithelium that terminally differentiates. Such complexity requires tightly orchestrated signals to balance differentiation with proliferation. We show here that skin commensals produce signals that directly regulate epithelial stratification through the xenobiotic sensor AHR. Future studies will be required to address the identity of the microbial metabolites that interact spatially within the complex architecture of the skin.

Our studies focused on the keratinocyte AHR and its role in forming the EPB, but the AHR can also be expressed by epidermal Langerhans cells, innate and adaptive immune cells, and dermal cells. Previous studies suggest that repair of the EPB is not dependent on AHR derived from Langerhans cells [58]. However, during epicutaneous sensitization, Langerhans specific loss of AHR led to decreased Langerhans cells number and function and dysregulated T cell responses [68]. Thus, AHR likely represents a sensor for Langerhans cell activation as part of the immunological barrier. Further studies will need to better define the cell-type specificity of AHR signaling to the different levels of barrier function, including the immune barrier.

Through depletion models and topical association with defined microbial consortia, we demonstrate the necessity and sufficiency, respectively, of commensal microbes in epithelial barrier function. We note that while colonization of germ-free mice with commensal microbes for two-weeks (COL) resulted in restoration of key epithelial differentiation signals, there were still differences between gene expression profiles of COL mice and SPF mice raised in the presence of microbes. It has been shown that skin-resident immune cells rely on imprinting by early life bacterial exposures to regulate acute wound healing in adult mouse skin [69]. Therefore, it is possible that certain skin development genes rely on early-life exposure to the microbiota for complete restoration of epithelial gene expression profiles.

Our data supports the hypothesis that sensing of microbial signals by the xenobiotic receptor AHR are crucial for self-renewal required of the epidermis. Topical application of coal tar is one of the oldest therapies for atopic dermatitis and has been shown to activate AHR to induce epithelial differentiation [70]. Recently, the natural product derived small molecule, Tapinarof, was found to bind and activate the AHR to moderate inflammatory responses in atopic dermatitis and psoriasis [71]. In support of the hypothesis that AHR mediates microbial signals to promote barrier function, we found that treatment with the potent and selective AHR ligand FICZ at a low dose was able to restore epithelial barrier repair in GF mice and induced epithelial differentiation. However, the role of AHR in skin barrier regulation may be highly context dependent. In murine models, exposure to pollutants can lead to hyperactivation of AHR that results in skin barrier damage and inflammation, which mirrors the phenotype of mice that constitutively express AHR in the keratinocyte [72, 73]. Thus, the balance in the specificity and quantity of AHR ligand, from endogenous and environment sources, is likely a key factor in modulating downstream signaling and impact on the skin, and requires further investigation.

AHR has now been recognized as an intracellular pattern recognition receptor that can identify and metabolize bacterial pigmented virulence factors, and promote antibacterial defense responses [74]. Our studies show that loss of AHR in skin led to increased susceptibility to the skin pathogen *S. aureus*. It remains to be determined whether increased susceptibility was due to impaired physical barrier, impaired antimicrobial barrier, or both. For example, the microbiota-induced antimicrobial protein RELMα protects against skin infection in a Vitamin A dependent manner [75]. Since there is interaction between AHR and retinoic acid signaling pathways [76, 77], this may represent a mechanism by which the skin microbiota mediates the antimicrobial barrier. The differential roles for pathogens and commensals in regulating AHR and promoting downstream effects are undefined, though are critical when considering diseases of barrier impairment or wounding, which are often complicated by *S. aureus* colonization and/or infection.

In summary, our findings show a role for skin microbiota in regulating epithelial differentiation and barrier function in stratified epithelia through AHR. These studies show that skin microbiome directly impacts development of the epidermal physical barrier. Future studies that address how microbial communities interact with each other to influence xenobiotic signals in homeostatic versus disease states will help leverage how personalized microbiota-based therapies can be used to improve the skin barrier.

## MATERIALS AND METHODS

### RESOURCE AVAILABILITY

#### Lead contact

Further information and requests for resources and reagents should be directed to and will be fulfilled by the corresponding author, Elizabeth Grice.

### EXPERIMENTAL MODEL AND SUBJECT DETAILS

#### Animal models and husbandry conditions

All mouse experiments were conducted under protocols approved by the University of Pennsylvania Institutional Animal Care and Use Committee (Protocol #804065). Age-matched 6-8 weeks old mice were used for all experiments. The following strains of mice were used in these studies: C57BL6/J (JAX stock #000664), Rag1 KO (JAX stock #002216)[78], Ahr^fx^ (JAX stock #006203)[79] and K14cre (JAX stock #018964)[80].

*<u>Germ-free studies</u>* were conducted in the Penn Gnotobiotic Mouse facility. Indicate strains were bred and maintained as germ-free (GF) in flexible vinyl isolators at the Penn Gnotobiotic Mouse facility housed in the University of Pennsylvania, School of Veterinary Medicine. Mice were housed as 3-5 mice per cage, until they were euthanized for tissue harvest. Aggressive mice or those that showed scratching wounds were not used in the studies. SPF counterparts were purchased from Jackson Laboratories and allowed to acclimatize in the facility for one week prior to beginning any experiments. One week prior to beginning of an experiment, mice (GF or SPF) were transferred to hermetically-sealed cages with individually filtered-positive airflow. The mice were maintained in these cages for the duration of the study. This allowed similar housing conditions for both GF and SPF mice for consistent TEWL readings. All mice were given autoclaved bedding, water and irradiated chow (5021 Autoclavable Mouse Breeder Diet, LabDiet®). Mouse handling was conducted in a laminar flow cabinet through double layer protective gloves. Germ-free status of mice was confirmed by weekly bacterial checks by the germ-free facility. At the end of each experiment, mice were autopsied and germ-free status was confirmed by enlarged cecum. Additionally, skin swabs, fecal pellet, and bedding samples were cultured by on blood-agar plates.

*<u>Studies in wild-type and AHR KO mice</u>* were conducted in Clinical Research Building vivaria at the University of Pennsylvania. C57BL6/J mice were maintained and bred by lab personnel. To generate *K14*^*Cre/+*^*Ahr*^*f/f*^ mice *Ahr*^*f/f*^ mice were crossed with *K14*^*Cre/+*^ mice to generate *K14*^*Cre/+*^*Ahr*^*f/+*^ F1 mice, *K14*^*Cre/+*^*Ahr*^*f/+*^ F1 mice were backcrossed to *Ahr*^*f/f*^ mice to obtain experimental mice (*K14*^*Cre/+*^*Ahr*^*f/f*^) and litter-mate controls (*Ahr*^*f/f*^) lacking *Cre*. To ensure robustness of studies, mice were randomly housed. Genotyping protocol: *Ahr*^*f/f*^ status was determined using PCR primers oIMR6075-Reverse (5’ CAG TGG GAA TAA GGC AAG AGT GA 3’) and oIMR6076-Forward (5’ GGT ACA AGT GCA CAT GCC TGC 3’) and resolving on a 5% polyacrylamide gel. *Cre* allele was determined using Generic Cre protocol (Protocol #22392, Jax Lab, Version 1.3) and resolved on a 2.5% Agarose gel.

#### Primary keratinocytes from adult mouse skin

Keratinocytes were derived from mouse tail or ear skin as described previously with slight modifications [81]. Following euthanasia mouse ears and/or tail were excised. With the help of forceps, ears were split into dorsal and ventral halves. To peel the tail skin from the bone, a scalpel was used to cut along the ventral axis from base of tail to tip. The exposed tail bone was peeled off using blunt-tip forceps. The resultant skin was cut into 0.75cm^2^ pieces. Skin obtained from ears and tails were incubated dermis side down in 6-well dishes and floated in ice-cold dispase (1mg/ml) in 1X PBS overnight at 4°C. Epidermal sheets were separated by lifting the epidermis using forceps. The separated epidermal sheets were cut into tiny pieces using forceps and scissors and incubated in 60mm untreated culture dish containing 2ml of 0.25% Trypsin-EDTA at 37°C, 5% CO2 for 15 minutes. At this point, 5ml suspension media (DMEM+10% FBS+ P/S) was added to the dish and the skin pieces were pipetted vigorously using a 10ml pipette to obtain a single cell suspension. Cell suspension was centrifuged at 150g for 5 minutes at 4°C and supernatant was removed. The cell pellet was suspended in 10ml suspension media and passed through 100µm cell strainer. Cell suspension was centrifuged at 150g for 15 minutes at 4°C, supernatant was removed and cells were suspended in 1ml suspension media and plated in collagen-coated 60mm dishes at 0.5 mouse equivalents (i.e. 5 million cells/ml) in plating media (low Ca2^+^ KSFM+ growth supplements+ 5% dialyzed FBS+4% DMEM) containing 10µM ROCK inhibitor (abcam120129) to prevent differentiation as described previously[82]. Typical cell counts were 2-5 × 10^6^ cells per mouse (cell count) and viability greater than 70% (cell viability) as determined by Trypan blue exclusion assay. Dishes were collagen coated by incubating 1.5ml collagen solution (50µg/ml collagen in 0.02N acetic acid) for 1 hour at 37 °C, 5% CO2 or overnight at room temperature. After incubation plates were rinsed 3 times with 1X PBS. Cells were remained undisturbed for 48 hours and then passaged for different experiments. Cells were maintained at 37°C in an atmosphere of 5% CO2 with humidity.

#### Human keratinocyte cultures

<u>Primary cultures of human keratinocytes</u> were obtained from neonatal foreskins through the Penn Dermatology Skin Biology and Diseases Resource-based Center: Skin Translational Research Core (STaR) Core B (visit: https://dermatology.upenn.edu/sbdrc/core-b/). Each experiment was conducted with at least three donors (as indicated in text). All experiments were conducted with cells at passage number less than 4. Briefly, cell suspensions were generated using dispase and trypsin, and the cells were cultured in a keratinocyte growth media [50% Medium 154, M154500 (Life Technologies), 50% Keratinocyte SFM, 17005042 (Life Technologies), 1% HKGS supplement, S0015 (Life Technologies), 1% Antibiotic/Antimycotic, 15240062 (Invitrogen)]. For routine passaging, cells were split when they were less than 70% confluent. Cells were washed with 1X PBS and trypsinized with 0.25% Trypsin-EDTA for 5 minutes, trypsin was inactivated using trypsin inhibitor (R007100, Thermo Fisher Scientific) and cell suspension was centrifuged. Following removal of supernatant, cell pellet was suspended in culture media and seeded as per experimental design. Cells were maintained at 37°C in an atmosphere of 5% CO2 with humidity.

<u>Immortalized human keratinocyte HaCaT</u>cells [83] were used for AHR reporter assay. HaCaT cells were verified for lack of mycoplasma contamination by ATCC. Experiments were conducted on cells that were at passage numbers between 26-36. For routine cell culture, HaCaTs were maintained in DMEM high glucose (11965092, Thermo Fisher Scientific) supplemented with 1% Sodium Pyruvate, 5% FBS, 1% Antibiotic/Antimycotic, 15240062 (Invitrogen) and 1% Non-essential amino acids). For routine passaging, cells were split when they were less than 70% confluent. Cells were washed with 1X PBS and trypsinized with 0.25% Trypsin-EDTA for 5 minutes, trypsin was quenched with DMEM containing 5% FBS. Cells were maintained at 37°C in an atmosphere of 5% CO2 with humidity.

#### Microbial strains

The <u>Flowers’ Flora Consortium,</u> consisted of *Staphylococcus epidermidis* (EGM 2-01), *Staphylococcus hemolyticus* (EGM 2-08), *Staphylococcus warneri* (EGM 2-09), *Micrococcus luteus* (EGM 2-04) and *Corynebacterium aurimucosum* (EGM 2-02), that had been isolated from healthy human skin and maintained in the Grice lab culture repository (number in parenthesis indicates identifier code in Grice lab culture collection). The *<u>S.aureus</u>* strain (AH3926) used in these studies was generously provided by Dr. Alexander Horswill (University of Colorado, Anschutz Medical Campus). *S. aureus* AH3926 consists of tdTomato stably integrated into *S. aureus* LAC (AH1263) and it’s construction has been described in detail previously [60]. Culturing conditions: All strains were cultured on solid blood agar plates at room temperature for 24-48 hours. For liquid cultures, all species (except *M. luteus*) were inoculated in tryptic soy broth and grown by shaking at 100rpm at 37°C. *M. luteus* was inoculated in nutrient broth.

## METHOD DETAILS

### RNA-sequencing of murine epithelia and analysis

Mice were shaved, and skin was collected from dorsal region. The fat layer was scraped off using a scalpel and then the skin was floated in dispase (1mg/ml) in 1X PBS overnight at 37°C for 1 hour in order to separate the epidermis from the dermis. The epidermis was stored in RNAlater. Mouse epidermis that had been stored in RNA-Later (Thermo-Fisher) was blotted dry and approximately 20 mg of tissue was placed in a Lysing Matrix A tube (MP Bio) with 600 µl RLT buffer (Qiagen) containing 2-mercaptoethanol. The tissue was homogenized with three, 1 min bursts of bead beating in a FastPrep 24 (MP Bio). The lysate was centrifuged (14,000 x g, 3 min) and the supernatant was transferred to a new tube to which 1 volume of 70 % ethanol was added. RNA was purified using a RNeasy Tissue Kit (Qiagen), as per manufacturer’s guidelines. RNA was quantified on a Qubit and RNA-integrity was assessed using BioAnalyser according to manufacturer’s instructions. 1µg RNA was used to construct RNA-seq libraries using the stranded-TruSeq RNA Sample Prep Kit (Illumina), spiked with phiX and sequenced on the Illumina NextSeq-500 Platform in 3 runs of 1×75 reads. The three runs were aggregated and then analyzed and aligned against the mouse genome [Genome Reference Consortium Mouse Build 38 patch release 5 (GRCm38.p5)] using AlignerBoost [84] and STAR 2.5.3[85]. Gene counts were fitted into a negative binomial model where both the gnotobiotic condition (SPF, GF or COL) and sex of the mouse were included using the DESeq2 [86] R package. Pairwise DEGs between conditions were obtained by setting corresponding “contrasts” for each pairwise comparison and filtering genes with FDP adjusted p-values less than 0.1. To identify enriched Gene Ontology (GO) terms, all annotated GO terms for aforementioned DEGs were retrieved using the ENSEMBL biomaRt R package[87], and significant enriched GO terms were identified using the topGO R package with the FDR-adjusted *p-*values < 0.1 under dependency [88].Uniquely enriched GO terms were selected by grouping similar GO terms using the online GO visualization tool REVIGO[89] with default (medium) similarity settings.

### 16S rRNA Gene Sequencing

#### Sample collection

Mice were swabbed prior to shaving with sterile foam-tipped applicators (Puritan) as described previously [49]. The swabs were snap frozen and stored at −80°C immediately following collection. Bacterial DNA was extracted from swabs as described [90]. In brief, swabs were incubated for one hour at 37°C with shaking in 300µL yeast cell lysis solution (from Epicentre MasterPure Yeast DNA Purification kit) and 10,000 units of ReadyLyse Lysozyme solution (Epicentre). Samples were subjected to bead beating for ten minutes at maximum speed on a vortex mixer with 0.5 mm glass beads (MoBio), followed by a 30-minute incubation at 65°C with shaking. Protein precipitation reagent (Epicentre) was added and samples were spun at maximum speed. The supernatant was removed, mixed with isopropanol and applied to a column from the PureLink Genomic DNA Mini Kit (Invitrogen). Instructions for the Invitrogen PureLink kit were followed exactly, and DNA was eluted in 50 mL elution buffer (Invitrogen). At each sampling event, swab control samples that never came into contact with the skin were collected, prepared and sequenced exactly as the experimental samples. No significant background contamination from either reagent and/or collection procedures was recovered.

#### Sequencing and analysis

Amplification of the 16S rRNA gene V1–V3 region was performed as described previously [90]. Sequencing was performed at the PennCHOP microbiome core on the Illumina MiSeq using 300 bp paired-end chemistry. The mock community control (MCC; obtained from BEI Resources, NIAID, NIH as part of the Human Microbiome Project: Genomic DNA from Microbial Mock Community B (Even, Low Concentration), v5.1L, for 16S rRNA Gene Sequencing, HM-782D) was sequenced in parallel. Sequencing of the V1-V3 region was performed using 300 bp paired-end chemistry. Sequences were preprocessed and quality filtered prior to analysis, including size filtering to 460-600 nucleotides. HmmUFOtu was used for sequence alignment and phylogeny-based OTU clustering as described previously [91]. Statistical analysis and visualization was performed using the phyloseq package [92] in the R statistical computing environment.

### Barrier recovery

Barrier was assessed as described [93, 94] by using noninvasive probe (Courage+Khazaka, Cologne, Germany) to measure transepidermal water loss (TEWL) by diffusion (Tewameter ®, TM300) according to the manufacturer’s instructions. The dorsal flanks of mice were shaved 24 hours prior to beginning of barrier analyses. Basal epidermal permeability barrier function was assessed 24 hours after shaving. Barrier was disrupted by tape stripping (3M Scotch High Performance Packaging Tape, 2”X800”) to achieve comparable insults between experimental and control groups as indicated for each experiment. Mice were anesthetized using isoflurane during TEWL measurements. TEWL measurements were averaged at 1-second intervals for a 30 second period. Indoor ambient temperature and mean relative humidity were recorded for each experiment. For barrier recovery assessment, TEWL was measured by placing probe at the same location on the dorsal flank of the mouse each time. For consistency, the same person made all TEWL measurements.

### Antibiotic treatment of mice

An antibiotic cocktail consisting of Metronidazole (1g/L), Sulfamethoxazole (0.8g/L), Trimethoprim (0.16g/L), Cephalexin (4g/L) and Baytril (0.025g/L) dissolved in drinking water containing Splenda (1 packet/250ml) as sweetener was provided to the mice for two weeks. To ensure decreased microbial burden, cages were changed 3 times a week for antibiotic treated mice as described previously [49]. Control cages were changed once a week, as per conventional policies to ensure microbial biodiversity.

### Differentiation assay

Confluent cells were trypsinized (0.25% Trypsin EDTA) and plated at 10^4^ cells/well in 12 well collagen coated dishes without ROCK inhibitor. Cells were allowed to grow to confluency (typically 2-3 days) and then media was switched to cornification medium i.e. traditional E-media without EGF (3 parts DMEM+ 1 part DMEM/F12 + 5% FBS + cholera toxin + insulin + adenine + hydrocortizone + antibiotics) for three days, to induce polarization and transmembrane junction formation in keratinocytes [95]. At the end of three days cells were scraped off and processed for qPCR analyses.

### Transepithelial electrical resistance (TEER) measurements

Keratinocytes were trypsinized (0.25% Trypsin EDTA) and plated at 10^4^ cells/well on collagen coated 12mm transwells with 0.4µm pore (Sigma-Aldrich; CLS3460). Twenty-four hours post-plating cells were 100% confluent and media was switched to KSFM containing 5% FBS and 1.6mM Ca^2+^. Every 24 hours, transepithelial electrical resistance (TEER) was measured using an epithelial volt/ohm meter (EVOM) using a STX2 manual electrode. To measure TEER, three readings were taken per chamber to cover three different areas of the transwell membrane longitudinally over three days. The electrode was cleaned with 0.5% bleach followed by 70% ethanol between each transwell. Readings are reported in ohms-cm^2^. *<u>FICZ and AHR inhibitor treatments:</u>* For experiments described in Figure 4, primary human keratinocytes were seeded on transwells and grown in presence of 100nM each of AHR ligand 6-formylindolo[3,2-b] carbazole (FICZ) (Sigma-Aldrich, #SML1489) and/or AHR inhibitor (Sigma Aldrich, #CH-223191) throughout the course of the experiments.

### Tissue preparation and immunofluorescence analysis

Murine skin tissue was collected and fixed in 4% paraformaldehyde and embedded in paraffin and sectioned at 6µm, as described previously[10]. Immunofluorescence protocols are described in detail at dx.doi.org/10.17504/protocols.io.k95cz86 [96]. Briefly, tissue sections were deparaffinized with xylenes and rehydrated with graded ethanol. Heat-induced antigen retrieval was performed in 0.01M citrate buffer, pH 6.0 and blocked in 10% normal goat serum. Antibodies against the following proteins were used at indicated dilution: Cytokeratin-10 1:1000 (Biolegend, #905401), Desmoglein 1a 1:200 (Abcam, #ab124798), Loricrin 1:500 (Biolegend, #905101). Alexafluor 594 conjugated goat-anti rabbit 1:1000 (ThermoFisher Scientific, #A32740) was used as secondary antibodies. Tissue was counterstained with Hoechst stain. Wide-field fluorescent images were acquired using by means of a 20X lens objective on a Leica DM6000 Widefield Fluorescence Microscope at the University of Pennsylvania, School of Veterinary Medicine Imaging core. For purpose of quantification 10-12 random images were taken in a blinded fashion at constant light exposure of 100 miliseconds for the Alexafluor 594 channel and 10 seconds for the Hoechst Stain. Images were processed using ImageJ software version 10.2 (NIH, Bethesda, MD). Krt10, Dsg1a, and Loricrin levels of each image were calculated by the integrated density of the signal, normalized to the Hoechst stain signal from the same area. For statistical analysis, each stain was calculated by taking average levels in each corresponding to 10-12 images per mouse and a two-sided T-test was used to determine the significance of signal differences between groups.

### Electron Microscopy

The ultrastructural analysis by EM was performed as described previously [97] at the VA Medical Center and Department of Dermatology at the University of California, San Francisco, United States. Skin samples were fixed in 2% glutaraldehyde and 2% paraformaldehyde and post-fixed in reduced ruthenium tetroxide before epoxy embedding. The samples were cut on a Leica Ultracut E microtome (Leica microsystems, Wetzlar, Germany) and imaged on a JEOL 100CX transmission electron microscope (JEOL, Tokyo, Japan) using a Gatan digital camera. For quantification, the thickness of the cornified envelope was measured in at least 25 randomly selected positions in 5 random high-powered electron micrographs of the mid stratum corneum from three mice of each colonization state. The observer recording these measurements was blinded to the groups.

## AHR reporter assay

Immortalized human keratinocyte HaCaT cells [83] were seeded in 96 well plates at 10^4^ cells/well in 100µl KSFM media supplemented with supplements. Twenty-four hours later, when cells were 80-90% confluent, each well was co-transfected with 20ng of *Renilla* luciferase DNA pGL4.74 (Promega) and 180ng of *Firefly* luciferase reporter plasmids: pGL4.23 which has xenobiotic response element (XRE) corresponding to *Cyp1A1* activity. Construction of plasmid is described previously[20]. Transient transfections of HaCaT cells was performed with Fugene® HD (Promega, #E2311) at 3:1 Fugene transfection reagent: DNA ratio according to manufacturer instructions. Twelve hours post-transfection, media containing transfection complexes was removed and replaced with media containing either indicated bacterial strains to represent MOI=0.1, 1 and 10, peptidoglycan (10ng) or FICZ (10nM). Twenty-four hours post-infection, dual luciferase readings (*Renilla* and *Firefly*) were read using Dual-Luciferase® reporter assay system (Promega, #E1910) on the BioTek Synergy HT fluorescence plate reader. Background correction was performed by subtracting readings of empty wells from observed readings and *Firefly* luciferase activity was normalized to *Renilla* luciferase. Relative response ratio to compare *CYP1A1* induction by each bacterial species to treatment with known AHR ligand FICZ (10nM) was computed as follows: RRR= [(experimental sample ratio)-(negative control ratio)]/ [(positive control ratio)-(negative control ratio)][98]. Experimental sample refers to cells treated with indicated bacteria or peptidoglycan; negative control refers to unstimulated cells and positive control refers to cells stimulated with 10nM FICZ.

### cDNA synthesis for qPCR analyses

Mouse skin or epidermis that had been stored in RNA-Later (Thermo-Fisher) was blotted dry and approximately 20 mg of tissue was placed in a Lysing Matrix A tube (MP Bio) with 600 µl RLT buffer (Qiagen) containing 2-mercaptoethanol. The tissue was homogenized with three, 1 min bursts of bead beating in a FastPrep 24 (MP Bio). The lysate was centrifuged (14,000 x g, 3 min) and the supernatant was transferred to a new tube to which 1 volume of 70 % ethanol was added. RNA was purified using a RNeasy Tissue Kit (Qiagen). Traces of DNA were removed with DNase-1 and the RNA was stored at −80°C.RNA was quantified (Qubit) and 10 ng RNA was used as a template for Superscript III (Invitrogen) reverse transcription with random hexamer primers. Following treatment with RNase H, the cDNA was stored at −20°C.

### Colonizing mice with human skin commensals

For liquid cultures of human skin commensals, *S. epidermidis, S. warneri, S. hemolyticus* and *C. aurimucosum* were inoculated in tryptic soy broth (TSB) and *M. luteus* was inoculated in nutrient broth media, respectively and grown by shaking at 200rpm for 16 hours at 37°C. Cultures were centrifuged, media was removed and bacterial pellets were suspended in PBS to obtain 10^9^ CFU/ml. Equal amount of each bacteria (10^9^ CFU) was combined in 5ml PBS and inoculated in bedding of mouse cages daily for 2 weeks. For each inoculum, a fresh batch culture was grown overnight.

### Mouse treatment with AHR ligand FICZ

Mice were treated with a low dose (100µg/kg) of AHR ligand 6-Formylindolo[3,2-b] carbazole (FICZ) (Sigma-Aldrich, #SML1489), that has been shown to be sufficient to induce expression of *Cyp1a1*[54]. A stock solution (4.5mg/ml) of FICZ was made in DMSO and diluted in 50% Acetone. Mice were shaved 24 hours prior to treatment and FICZ was applied topically by directly pipetting onto shaved skin, daily for 2 weeks (**Figure 3B)**.

#### *S. aureus* skin infection protocol

##### Epicutaneous infection

Infection protocol for epicutaneous infection with *S. aureus* has been described previously [99] and was implemented with slight modifications. Briefly, mice were anesthetized with isoflurane, tape stripped (TEWL=20 g/m^2^/h) and 24 hours later, 10^7^ CFU *S. aureus* in 100µl was applied to back skin and spread using a swab. The inoculum was allowed to dry for 10 minutes and mice were returned to their cages.

##### CFU enumeration

Forty-eight hours post-infection, mice were euthanized and approximately 1cm^2^ infected skin area was collected and weighed and transferred to tubes containing 300µl 1X PBS. Tissue was homogenized by bead beating for twenty minutes at maximum speed on a vortex mixer with 0.5 mm ceramic beads and CFUs were enumerated by serial dilution on blood agar plates after overnight incubation at 37°C. Both *S. aureus* and total bacterial counts were determined and normalized to weight of tissues. *S. aureus* colonies were visible as red colonies due to stable expression of tDTomato and could be distinguished from total bacteria.

### Epicutaneous sensitization with ovalbumin

Procedures to induce barrier defects that mimic atopic dermatitis by repeated epicutaneous sensitization by ovalbumin (OVA) [30, 100] followed by infection with *S. aureus* [101] have been described previously. The dorsal skin of mice was shaved and TEWL was assessed to give a baseline reading. To measure and compare barrier repair between wild-type and AhR null mice, we standardized the amount of initial barrier disruption to give identical TEWL values. To achieve this, mice were tape stripped to give a reading of TEWL ∼20g/m^2^/h. Twenty-four hours post tape-stripping, the mice were treated daily for 7 days with 100 µg OVA (Sigma Aldrich, # A5503) suspended in 100 µl PBS was applied onto backs of mice and spread using a skin swab, and allowed to dry for 2 minutes. For inducing atopic dermatitis-like condition, the 7-day OVA treatment regime was repeated twice more, with 2 weeks rest between subsequent treatments. At the end of treatment mice were mice were tape stripped to give a reading of TEWL ∼40g/m^2^/h and *S. aureus* was applied as described earlier or barrier recovery was assessed.

## QUANTIFICATION AND STATISTICAL ANALYSIS

### Data visualization and statistics

All statistical analysis was performed using functions built into the R statistical environment (RStudio Version 1.3.1056). Data was visualized using ggplot2 [102] package and GraphPad Prism version 8.0.0 for Mac OS X, GraphPad Software, San Diego, California USA, www.graphpad.com. TEWL/TEER vs time readings were fitted by linear modeling function in R statistical package and visualized using ggplot2 package. Significance was assessed by ANCOVA analysis. Fit parameters (adjusted *R*^*2*^ and *F*-statistics) are indicated for each plot. Span indicated by shaded area represents 95% CI. Gene expression analysis from qPCR was conducted as per guidelines described previously[103]. Cycle thresholds were normalized to housekeeping genes (Rplp2, Sptbn1 and 18s rRNA) and normalized relative to quantitative cycle (Cq) values of control. Heatmaps for qPCR analysis were made using Morpheus heat map viewer from Broad Institute (https://software.broadinstitute.org/morpheus). Each square represents average normalized readings (n= 3 technical replicates). Bonferrroni correction was used to adjust for multiple comparisons. The AHR pathway was originally downloaded from wikipathways (https://www.wikipathways.org/index.php/Pathway:WP2873), then modified by highlighting DEGs identified in our analysis using a customized Perl script. The log2 fold-change values between SPF and GF mice were used to determine the color hue (red/blue: up/down in SPF vs. GF) and saturation of the highlighted DEGs.

## KEY RESOURCES TABLE

### SUPPLEMENTARY MATERIALS

**Table S1. Related to Figure 1**.

Results from differential expression analysis of RNA-seq data for genes depicted in Fig. 1F. As a result, we identified 6,396, 427, and 3,232 DEGs for SPF vs. GF, COL vs. GF and SPF vs. COL comparisons, respectively. DEGs defined as those with FDR adjusted p-values < 0.1. In Sheet SPF vs GF (upregulatedin SPF: log2FC> 0, downregulated in SPF: log2FC < 0); in sheet COLvsGF (upregulated in COL: log2FC> 0, downregulated in COL: log2FC< 0); in sheet SPFvsCOL (upregulated in SPF: log2FC > 0, downregulated in SPF: log2FC < 0).

**Table S2. Related to Figure 1**. Contains list of genes manually curated based on literature analyses that have been implicated in barrier function. Genes are listed under the following categories based on their functions: adherence junction formation, lipid-lamellae formation, keratin network, differentiation, skin barrier development, formation of cornified envelope and basement membrane.

**Table S3. Related to Figure 1**. Contains the key genes involved in skin barrier function that were differentially expressed between SPF and GF murine skin and were used to generate graph depicted in Figure 1F.

**Table S4. Related to Figure 1**. Contains list of differentially expressed barrier genes involved in skin barrier function that were differentially expressed between SPF and GF murine skin by using genes described in Table S2 as reference.

**Figure S1. Related to Figure 1**. Structural analysis of murine dorsal skin from SPF and GF mice. **(i)** Histopathological analysis of Hematoxylin and Eosin stained tissue did not demonstrate overt differences between age-matched GF and SPF mice. **(ii)** Electron Microscopy (EM) was performed on skin and number of layers of cornified envelope were counted. *** indicates *P<0*.*01*, T-test.

**Figure S2. Related to Figure 2**. Assessment of **(A)** TEWL and **(B)** pH at baseline in age-matched age-matched germ-free (GF) and specific pathogen-free (SPF) C57BL6/J mice (n=4 mice per group). **(C)** Shown is indicated TEWL corresponding to number of tape strips in GF vs SPF mice. **(D)** GF and SPF *Rag1−/−* mice were swabbed colony forming units (CFU) were determined and **(E)** TEWL was measured at baseline. **(F)** Dorsal skin from GF and SPF *Rag1−/−* mice was collected and gene expression was analyzed by qRT-PCR. Cycle thresholds were normalized to housekeeping genes (*Rplp2, Sptbn1* and *18s rRNA*) and normalized relative to Cq values of SPF condition. Each square represents average readings from three technical replicates from each mouse. \**P<0*.*01*, ** *P<0*.*001* by T-test adjusted by Bonferrroni correction. **(G)** Primary keratinocytes were derived from germ free mice and grown on transwells in basal medium before switching to cornification media containing 5% FBS and 1.6mM Ca^2+^. Shown here is expression of genes between GF and SPF mice before and after calcium switch. Expression of genes involved in differentiation [lnvolucrin *(Ivl)*, cytokeratin-10 *(Krt10)*] and adherence [Corneodesmosin *(Cdsn)*, Desmocollin-1 *(Dsc1)*, Desmoglein-1a *(Dsg1a)*] and AHR pathway (*Cyp1a1, Cyp1b1, Hsp90ab1*) was assessed by qRT-PCR. Cycle thresholds were normalized to housekeeping genes (*Rplp2, Sptbn1* and *18s rRNA*) and normalized relative to Cq values of SPF condition. Each square represents average readings from technical replicates (n= 4 technical replicates) derived from an individual mouse (n=4 mice per group). \**P<0*.*01*, ** *P<0*.*001* by T-test adjusted by Bonferrroni correction. **(H)** Primary keratinocytes grown on transwells were grown in 5% FBS and 1.6mM Ca^2+^ epithelial adhesion was assessed by measuring transepithelial electrical resistance (TEER) at indicated time points. Data from three individual experiments are shown. One dot represents average TEER readings from technical replicates (n=3) derived from one individual mouse (n=4 mice per group).

**Figure S3. Related to Figure 2**. To decrease skin microbial burden, wild-type SPF mice were treated with antibiotic cocktail (n=5) or vehicle (n=6) for two weeks. Genomic DNA was extracted from skin swabs and fecal samples collected at baseline (Day 0 i.e. D0) and after one week of treatment (Day 7 i.e. D7). V1-V3 region was amplified and analyzed by 16S rRNA gene sequencing. **(A)** Microbiota composition of each sample shown as the relative abundance of the top 15 most abundant genus of the entire dataset. **(B)** Fecal pellets collected at end of treatment (i.e. 14 days later) were weighed, homogenized and plated on blood agar plates and CFUs were enumerated. **(C)** Within-sample alpha diversity for samples from groups shown as the Shannon index. The within sample Shannon index was calculated using the Phyloseq R package. **(D)** Principle Coordinates Analysis (PCoA) plot of between microbiota sample beta-diversity for different sample types (skin or fecal) and antibiotic treatment (control or antibiotic). Ctrl: control treatment, Abx: Antibiotic treatment. Treatment timepoint was color labeled. The Bray-Curtis dissimilarity was used for the between-sample beta-diversity metric.

**Figure S4. Related to Figure 4**. TEWL dynamics during the course of ovalbumin sensitization treatment regime. **(A)** Schematic depicts sequence of manipulations to mice during OVA sensitization. Mice were tape stripped at the beginning of the experiment (TEWL∼20 g/m^2^/h) and OVA was applied daily for 7 days, 3 times with rest for 2 weeks between each treatment. At the end of final treatment, mice were tape-stripped (TEWL∼40 g/m^2^/h) and *S. aureus* was applied to back skin. **(B)** TEWL readings were taken at indicated time points in **(i)** *K14*^*Cre/+*^*Ahr*^*f/f*^ and **(ii)** *Ahr*^*f/f*^ mice treated with OVA or vehicle (PBS). Endpoint readings are reported in Figure 4. **(C)** Total bacterial counts per gram of tissue are reported for *K14*^*Cre/+*^*Ahr*^*f/f*^ (KO) and *Ahr*^*f/f*^ (WT) mice for indicated comparisons.

**Figure S5. Related to Figure 5**. TEWL dynamics during the course of short-term ovalbumin sensitization treatment regime after Flowers’ Flora colonization **(A)** Schematic depicts sequence of manipulations to C57BL6/J mice. Mice were pre-colonized with Flowers’ Flora daily for 2 weeks. At the end of pre-colonization, mice were tape stripped at the beginning of the experiment (TEWL∼20 g/m^2^/h) and OVA was applied daily for 7 days. At the end of final treatment, mice were tape-stripped (TEWL∼40 g/m^2^/h) and barrier recovery was assessed. **(B)** TEWL readings were taken at indicated time points in **(i)** mice pre-colonized with Flowers’ Flora and **(ii)** those treated with vehicle (PBS).

## ACKNOWLEDGEMENTS

We thank Dr. Alex Horswill (University of Colorado) for the kind gift of the *S. aureus* strain; Dr. Jorge Henao-Mejia (University of Pennsylvania) for the kind gift of *Ahr*^*f/f*^ mice; Dr. John Seykora (University of Pennsylvania) for the kind gift of HaCaT keratinocytes; Dmytro Kobuley, Gnotobiotic Mouse Facility (University of Pennsylvania) and current and former members of the Grice lab and the Department of Dermatology for critical discussion and review of the work.

