## Supplemental Figures for "Commensal Microbiota Regulates Skin Barrier Function And Repair Via Signaling Through The Aryl Hydrocarbon Receptor"

### Figure S1. Related to Figure 1.

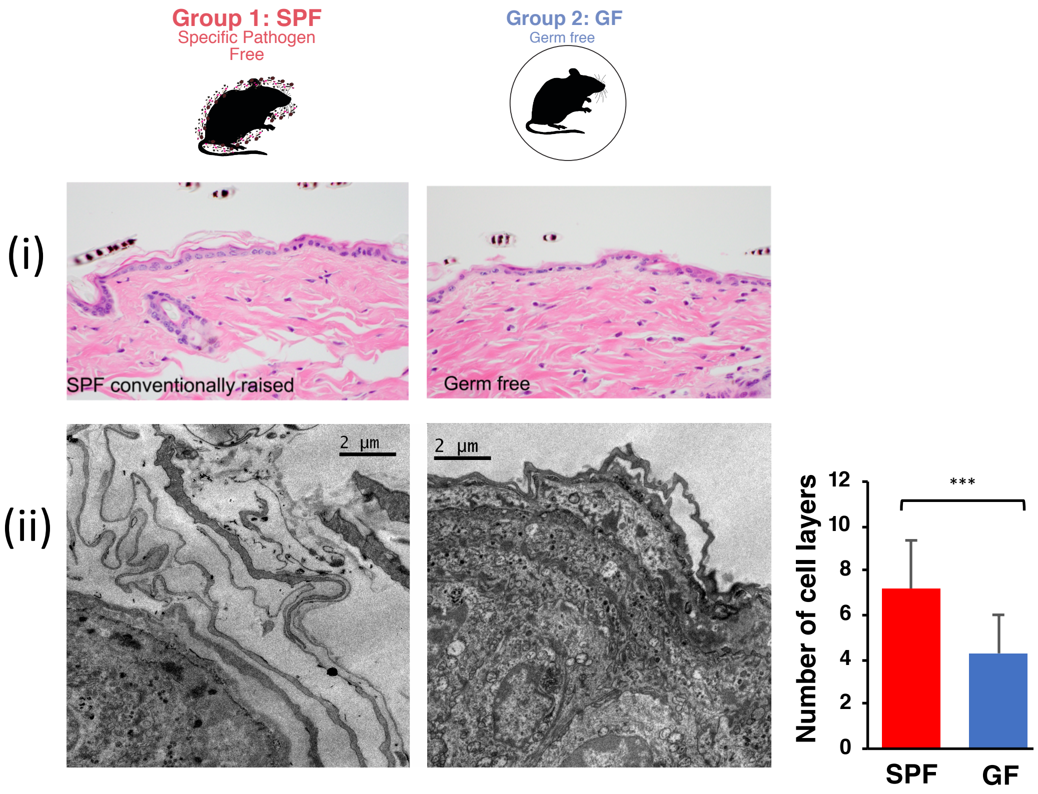

### Figure S2. Related to Fig. 2.

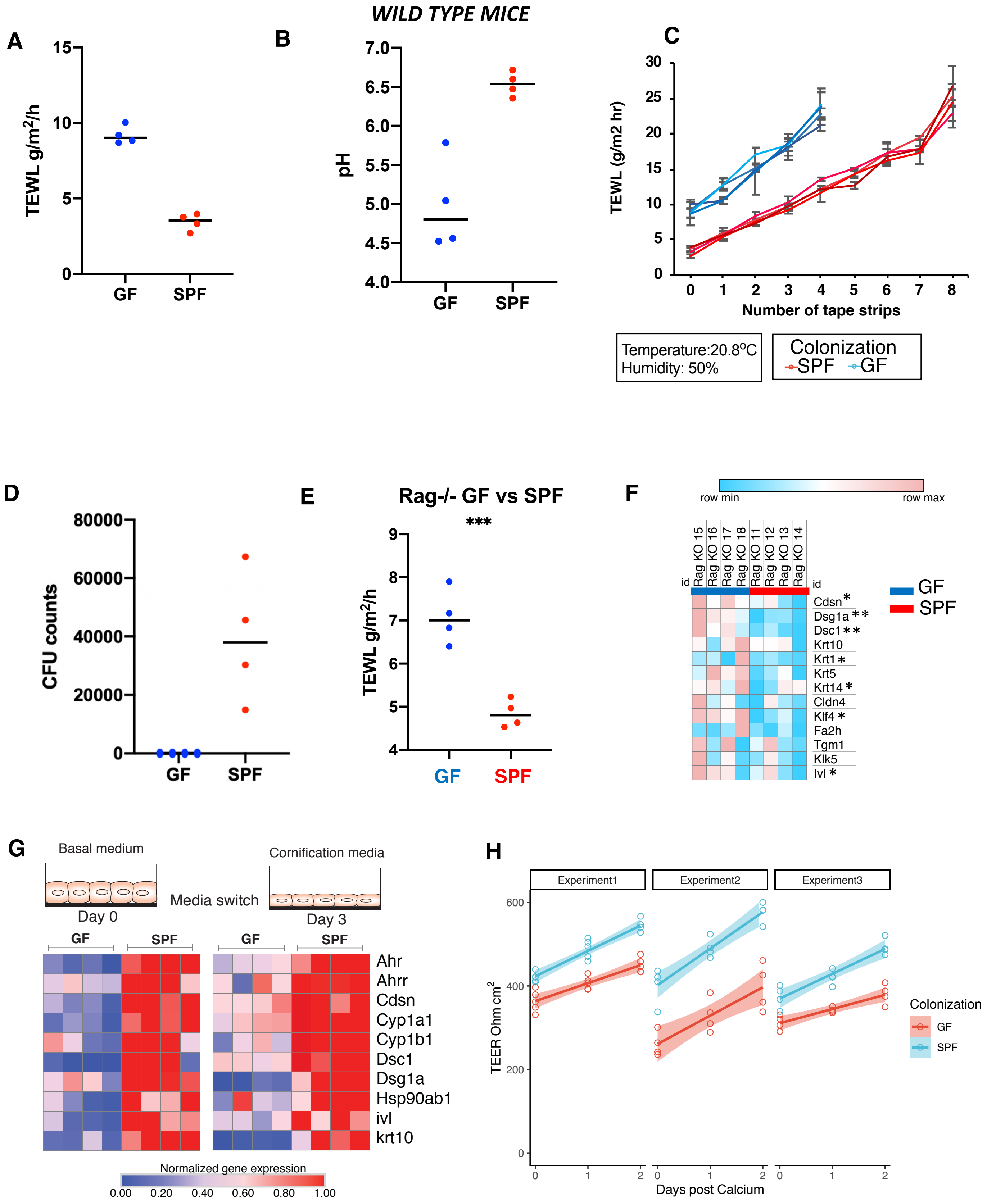

### Figure S3. Related to Fig. 2.

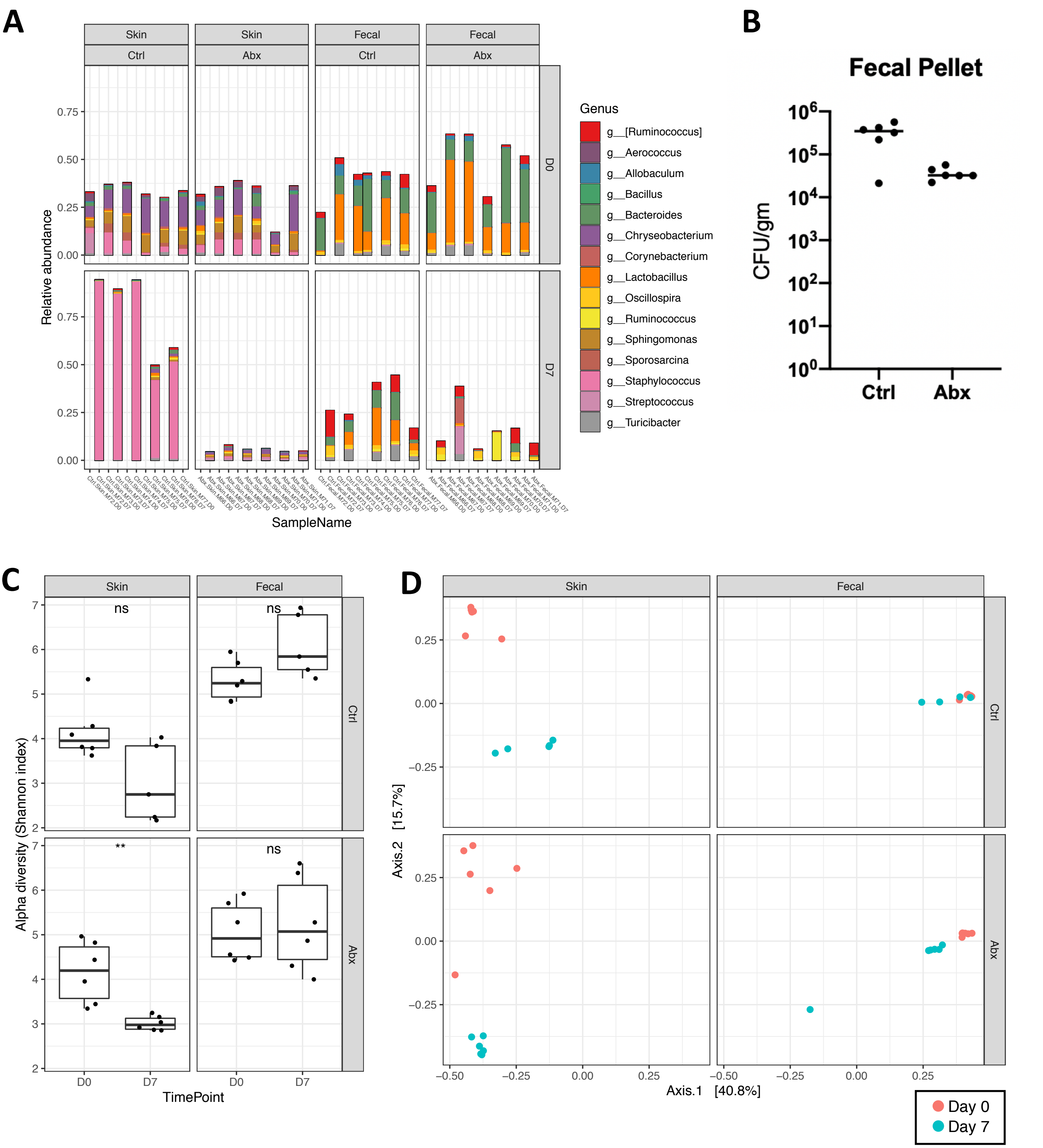

### Figure S4. Related to Figure 4

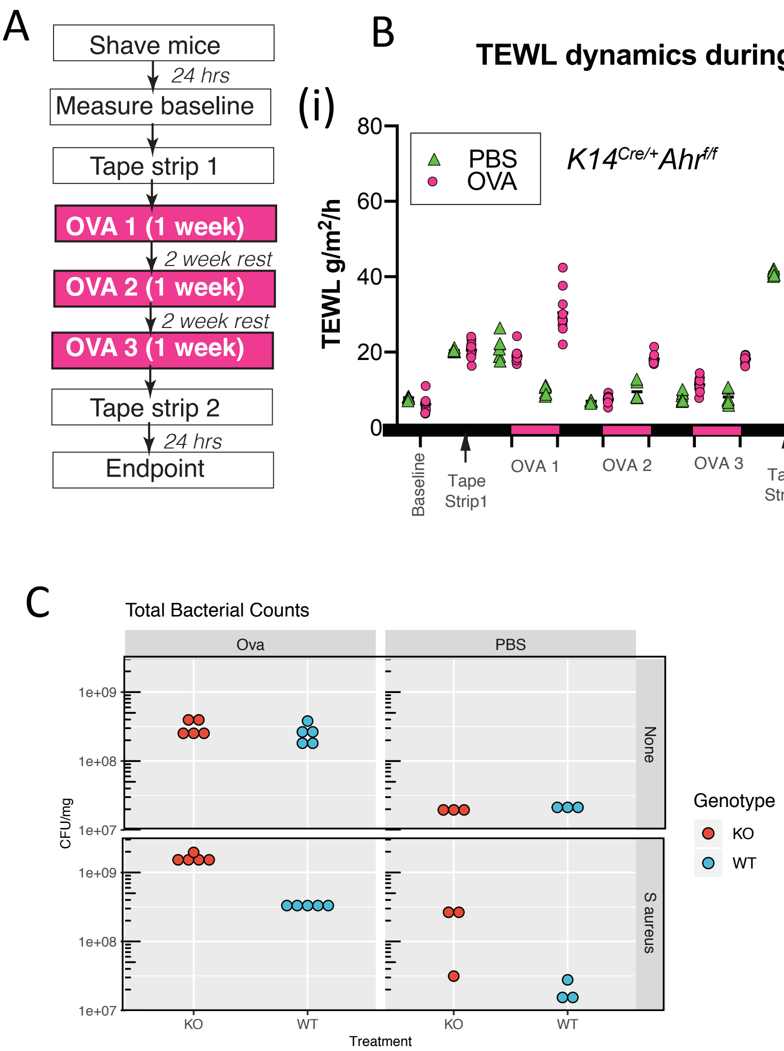

### Figure S5. Related to Figure 5

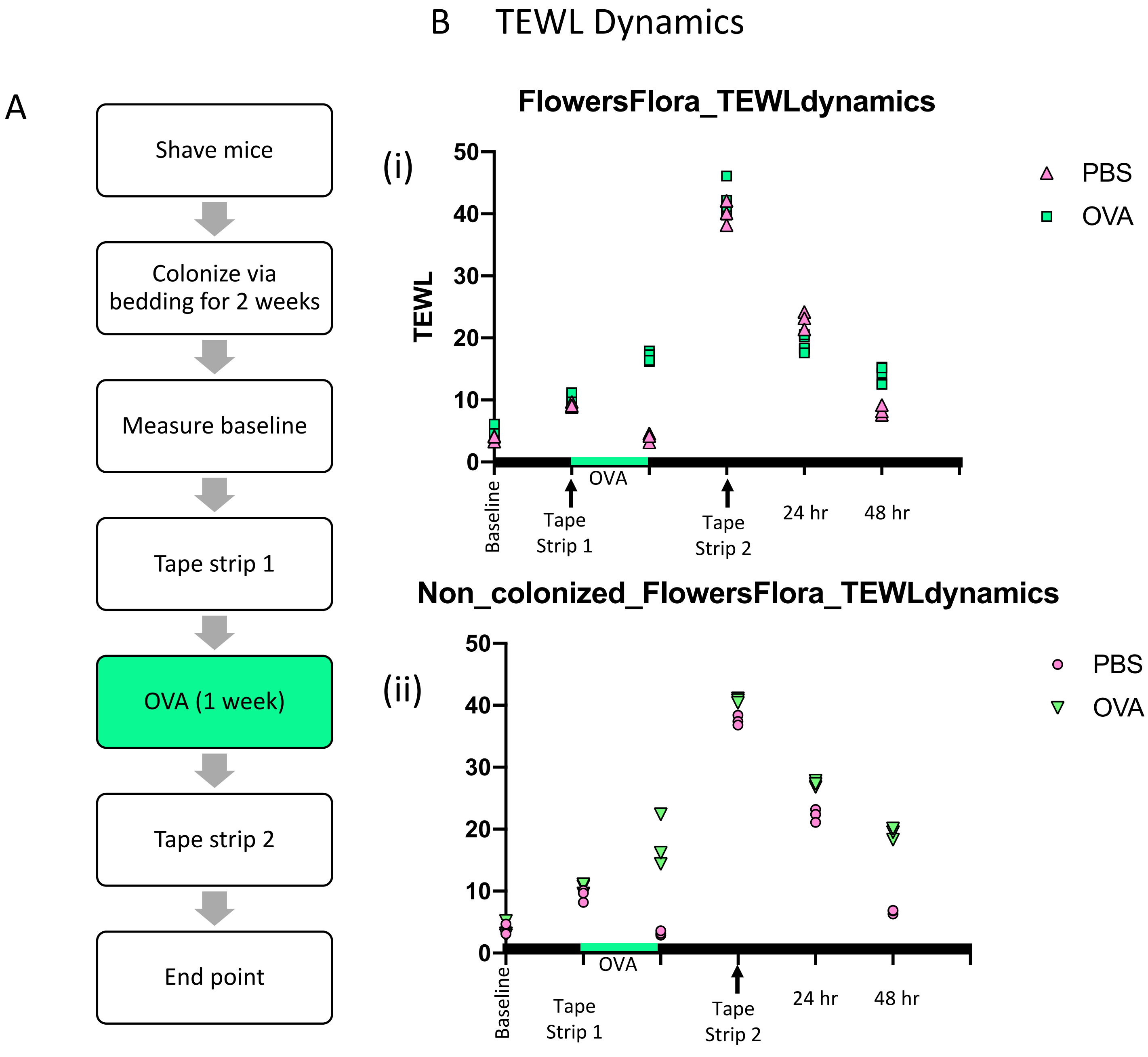
